# Aggregation-prone alpha-synuclein proteoforms and dysregulated molecular signatures in the vermiform appendix of synucleinopathy patients

**DOI:** 10.1101/2025.10.07.680938

**Authors:** Ehraz Anis, Saima Zameer, Joshua Wierenga, Peipei Li, Jacek W. Sikora, Juozas Gordevicius, Meghan Schilthuis, Richard D. LeDuc, Jeffery H. Kordower, Michelle Pinho, Sandra Pritzkow, Claudio Soto, Patrik Brundin, Lena Brundin, BA Killinger

## Abstract

Synucleinopathies, including Parkinson’s disease, are neurodegenerative diseases characterized by intracellular inclusions containing the amyloidogenic protein alpha-synuclein. While classically considered to be brain disorders, increasing evidence suggests involvement of the gut, with alpha-synuclein aggregates potentially propagating to the brain via the vagus nerve. Evidence also suggests that the vermiform appendix is particularly susceptible to alpha-synuclein aggregation, and appendectomy impacts the onset of Parkinson’s disease. However, the mechanisms underlying the aggregation of alpha-synuclein in the vermiform appendix remains poorly understood. To explore this, we assessed aggregation properties in postmortem appendix tissues from healthy controls and synucleinopathy patients using the alpha-synuclein seed amplification assay (alpha-synuclein-SAA) and performed total RNA sequencing alongside differential bisulfite-hybridization-based DNA methylation analysis in the same tissues to investigate the molecular underpinnings. Moreover, we determined alpha-synuclein cleavage patterns by cataloging soluble alpha-synuclein proteoforms from postmortem substantia nigra and post-surgical appendix tissues using top-down mass spectrometry (TD-MS). Alpha-synuclein-SAA was positive in appendix samples for 68.75% of synucleinopathy patients and 6.6% of controls. Genomic profiling revealed dysregulated expression of genes linked to protein folding/degradation, immune/inflammatory responses, and ciliary dynamics in synucleinopathy appendix tissues. TD-MS identified 65 distinct alpha-synuclein proteoforms in the substantia nigra and appendix, with 9 unique to the appendix. Further, *in silico* modeling revealed higher aggregation propensity of alpha-synuclein proteoforms in the appendix versus substantia nigra. Together, our findings suggest that a tissue environment of alpha-synuclein dysproteostasis in the appendix has the potential to contribute to the development of synucleinopathies.

**One Sentence Summary:** Appendixes from synucleinopathy patients show altered gene expression, unique α-syn proteoforms, and higher aggregation propensity than substantia nigra.

## INTRODUCTION

Synucleinopathies, including Lewy body diseases such as Parkinson’s disease (PD), are common neurodegenerative diseases characterized by accumulation of aggregated alpha-synuclein in the central and peripheral nervous systems. Alpha-synuclein is an intrinsically disordered protein that misfolds into beta-sheet-rich structures that are detergent-insoluble, protease resistant, and act via a prion-like aggregation mechanism (*1, 2*). The N-terminal domain (1-60 residues) folds into alpha-helical structures when interacting with lipid membranes (*3*) and the C-terminus (96-140 residues) remains largely disordered. Both the N- and C- terminus shield core residues, termed the non-amyloid component (NAC) domain (61-95 residues), which is required for aggregation.

Synucleinopathies are classically characterized by central nervous system (CNS) pathology, but increasing evidence highlights a role for alpha-synuclein in peripheral tissues. In particular, the gastrointestinal tract has been proposed as a potential origin site for PD pathology (*4–6*). Gastrointestinal dysfunction is a common non-motor symptoms of PD, with about 80% of patients reporting constipation years before the onset of motor symptoms (*7*). Moreover, misfolded alpha-synuclein is present in the enteric nervous system in PD, and is suggested to propagate from the gastrointestinal tract to the brain via the vagus nerve, among other potential neural pathways (*8*).

The vermiform appendix, a lymphoid-rich organ in the gastrointestinal tract that is highly innervated by enteric neurons, harbors monomeric and aggregated alpha-synuclein in both healthy individuals and in patients with PD (*9*). The presence of insoluble/aggregated alpha-synuclein in healthy appendix suggests that it does not necessarily have to trigger a synucleinopathy in the CNS; other factors likely contribute to the accumulation and propagation of pathogenic alpha-synuclein to the brain. Epidemiological findings have previously suggested that removing the appendix early in life might reduce the risk of PD and delay the onset of PD late in life (*9*). In another study, chronic appendicitis-like lesions were detected using multislice spiral computed tomography (MSSCT) in 53% of PD patients, compared to 4% in matched controls (*10*). This suggests that chronic inflammation of the appendix is associated with PD in a subset of patients.

Aberrant gene expression and epigenetic modifications have been reported to contribute to synucleinopathy through alterations in cellular homeostasis, stress response pathways, and compromising neuronal survival (*11, 12*). An earlier study from our team demonstrated epigenetic and gene expression abnormalities specifically in the autophagy-lysosomal pathway in the PD appendix (*13*). Additionally, perturbations in other proteostasis machinery responsible for the degradation of misfolded proteins, like the ubiquitin-proteasome system or chaperone-mediated autophagy (*14*), could contribute to the accumulation of aggregated alpha-synuclein in the appendix. Numerous studies have also consistently shown an association between inflammation and alpha-synuclein aggregation, although the underlying mechanisms remain to be fully elucidated (*15, 16*). Notably, growing evidence from clinical and pre-clinical studies supports a significant link between chronic gut inflammatory conditions like inflammatory bowel disease (IBD) and PD (*17, 18*).

Specific N- and C-terminus truncation enhances alpha-synuclein aggregation propensity and tendency to propagate from one neuron to another (*19*), provided the alpha-synuclein NAC domain (61-95 residues) remains intact. Several truncated proteoforms are enriched in Lewy pathology and show enhanced aggregation kinetics *in vitro* (*20–22*). Past efforts have been focused on identifying post-translational modifications (PTMs) of insoluble alpha-synuclein, and consequently, less is known about PTMs of the soluble alpha-synuclein pool in synucleinopathies. This is despite evidence that smaller soluble oligomeric structures are key players in toxicity (*23*) and pathology spread (*24*). Moreover, the truncation of alpha-synuclein in peripheral tissues and its functional significance in disease pathogenesis is poorly understood.

While previous research has established a link between the appendix and PD, the complex, multi-layered disease-related changes within the appendix that could potentially explain this link remain unclear. We have taken an integrated approach, combining alpha-synuclein-seed amplification assay (alpha-synuclein-SAA), transcriptomic (RNA sequencing), and epigenomic (DNA methylation profiling) analyses from the same appendix tissues, which allows for a holistic understanding of disease-associated molecular changes. Additionally, we comprehensively cataloged alpha-synuclein proteoforms in the post-surgical appendix and substantia nigra (SN) of patients with synucleinopathy and neurologically intact controls using top-down mass spectrometry (TD-MS), the gold-standard approach for identifying intact and truncated alpha-synuclein proteoforms (*25, 26*). Based on our previous findings and other recent literature linking alpha-synuclein aggregation propensity (*27*), aberrant gene regulation (*28*), and truncation (*29*) to propagation of pathology, we hypothesized that aberrant gene regulation in appendices of certain individuals facilitates accumulation of pathogenic alpha-synuclein and thereby increases risk of developing synucleinopathy.

## RESULTS

### Catalog of alpha-synuclein proteoforms in the human appendix and SN

To determine whether the appendix contains alpha-synuclein proteoforms relevant to disease pathology, we profiled the alpha-synuclein proteoforms in the appendix of synucleinopathy patients and healthy individuals and performed comparisons to those present in the SN. In total, 65 distinct alpha-synuclein proteoforms were identified in the human appendix (Figure 1A), ranging from the full-length N-terminally acetylated proteoform (alpha-synuclein^ac1-140^) to truncated species 31 amino acids in length (alpha-synuclein^78-109^). There were truncated species with N- or C-terminal cleavage only (n= 4 and 13 alpha-synuclein proteoforms, respectively). However, the large majority, 46 alpha-synuclein proteoforms, were cleaved at both ends: 23 with N- and C-terminal truncations and 23 with intra-NAC domain and C-terminal truncations. In total, we identified 27 unique cleavage locations for alpha-synuclein proteoforms in the appendix, of which 6 occurred within the N-terminus, 6 were in the NAC domain, and 15 were in the C-terminus. The most common truncation site was in the C-terminus at position 114, which has been found to promote Lewy pathology formation in neurons (*30*), and other frequently cleaved sites included positions 40, 73, 109 and 115 of alpha-synuclein (number of proteoforms with cleavage site: 13, 9, 9, 8, 8, respectively). Of the 65 alpha-synuclein proteoforms identified in the appendix, 56 were also detected in the SN. All proteoforms detected in the substantia nigra were present in the appendix. We found 9 alpha-synuclein proteoforms that were present only in the appendix, with most of these cleaved in both the NAC domain and C-terminus (alpha-synuclein^66-114^, alpha-synuclein^68-114^, alpha-synuclein^73-114^, alpha-synuclein^66-132^, alpha-synuclein^68-109^, alpha-synuclein^73-122^, and alpha-synuclein^73-125^). Thus, the appendix contained a diverse and distinct pool of alpha-synuclein truncation products.

**Figure 1.**
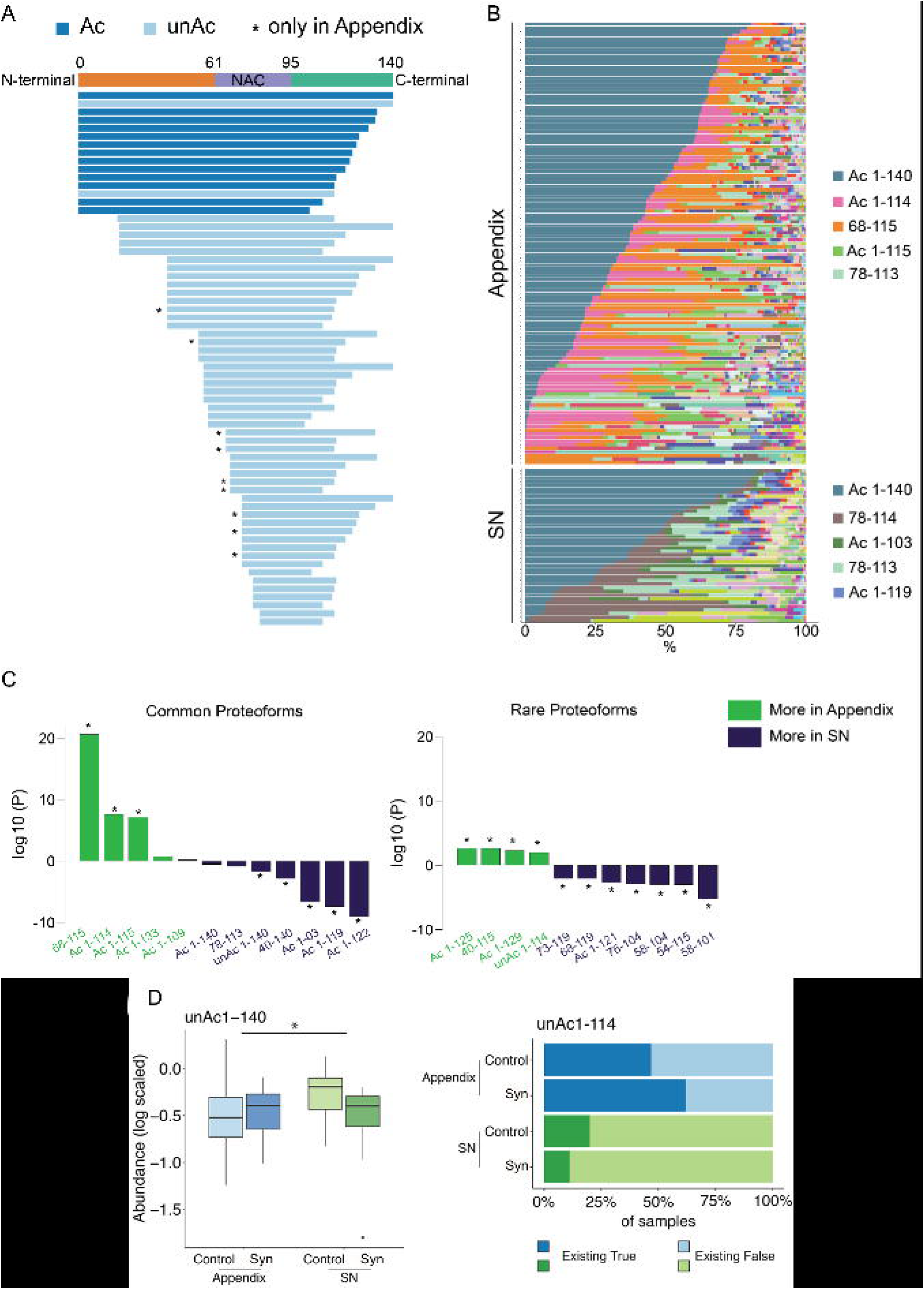
Alpha-synuclein proteoforms in the appendix and substantia nigra (SN) **(A)** Bar plot showing 65 alpha-synuclein proteoforms identified in the human appendix and SN. Acetylated proteoforms are colored dark blue; unacetylated proteoforms are colored light blue. Nine proteoforms, which were only found in the appendix, are denoted with *. **(B)** Bar plot showing alpha-synuclein proteoforms distribution according to their abundance in the human appendix and SN separately. X-axis shows the percentage of the abundance for each existing proteoform. Top 5 most abundant proteoforms are listed with their color codes. **(C)** Common alpha-synuclein and rare proteoforms showing tissue differences. For common alpha-synuclein proteoforms, robust linear regression was used for statistical analysis adjusting for diagnosis (synucleinopathy/control) and age, with proteoform abundance as dependent variable (Y) and tissue as independent variable (X). Proteoforms showing significant tissue differences after FDR q<0.05 are denoted with *. For rare alpha-synuclein proteoforms robust linear regression was used for statistical analysis adjusting for diagnosis (synucleinopathy/control) and age, with proteoform detection (TRUE or FALSE) as dependent variable (Y) and tissue as independent variable (X). Only rare proteoforms showing tissue difference after FDR q<0.05 are shown and denoted with *. (More in appendix in color dark green; more in SN in color dark blue). **(D)** unAc1-140 increased in the appendix, unAc1-114 increased in the SN. Robust linear regression was used for statistical analysis, adjusting for diagnosis (synucleinopathy/control) and age.

We next compared the abundance of alpha-synuclein proteoforms in the appendix and SN (Figure 1B, 1C). First, we determined whether the appendix and SN had quantitative differences in common proteoforms (present in ≥50 samples). We found that alpha-synuclein proteoforms with truncations at the C-terminus position 114/115 were significantly more common in the appendix than in the SN (*q*<0.05 for alpha-synuclein^68-115^, alpha-synuclein^1-114^ and alpha-synuclein^1-115^, linear regression). In contrast, the appendix had lower levels of other C-terminal truncations, including at positions 103, 119, and 122, relative to the SN (*q*<0.05 for alpha-synuclein^1-103^, alpha-synuclein^1-119^ and alpha-synuclein^1-122^, linear regression). To analyze the rare proteoforms, we determined whether each alpha-synuclein proteoform occurred more frequently in the appendix or SN (Fig. 1C). We found that in the appendix several rare proteoforms were more abundant (alpha-synuclein^ac1-125^, alpha-synuclein^40-115^, alpha-synuclein^ac1-129^, alpha-synuclein^ac1-114^) and conversely, several rare proteoforms were more abundant in the SN (alpha-synuclein^ac1-121^, alpha-synuclein^58-101^, alpha-synuclein^54-115^, alpha-synuclein^58-104^, alpha-synuclein^76-104^, alpha-synuclein^68-119^, alpha-synuclein^73-119^). We also found that levels of reliably detected proteoforms alpha-synuclein^ac1-122^ and alpha-synuclein^78-113^ were significantly depleted in synucleinopathy cases relative to controls (q<0.05, linear regression; Supplementary Figure S1). We also detected instances of syn proteoforms that were unacetylated at the N-terminus for alpha-synuclein^1-140^ and alpha-synuclein^1-114^, in addition to their abundant N-terminal acetylated counterparts in both the appendix and SN (Fig. 1D). Therefore, both common and rare proteoforms differed in relative abundance between appendix and SN. Through classifier model training, we predicted with an accuracy above 80% that alpha-synuclein^ac1-122^ may be a highly relevant predictor of synucleinopathy in both appendix and SN (Supplementary Figure S1).

We further investigated age-dependent changes in the alpha-synuclein proteoforms present in the appendix and SN and found that synucleinopathy appendix had a significant gain in full-length synuclein (alpha-synuclein^ac1-140^) with age, which was not observed in the appendix of unaffected controls (q<0.05, linear regression; Supplementary Figure S2). Finally, sample-wise Pearson correlation showed that synuclein proteoforms in the appendix are more similar to those in synucleinopathy SN than control SN (Supplementary Figure S3).

### The aggregation propensity of alpha-synuclein in the appendix

Given that alpha-synuclein aggregation plays a central role in the development of synucleinopathies, we sought to characterize the aggregation propensity of the proteoforms we identified in the appendix and SN. We modelled the aggregation potential of each alpha-syuclein proteoform using a computational approach that determines the energy requirements for amyloid fibril formation (*31*). We then determined the alpha-synuclein aggregation propensity score for each sample based on proteoform abundances and their aggregation potential. We found that the aggregation propensity of alpha-synuclein in the appendix was significantly higher than in the SN (*p*<10^-10^, linear regression; Figure 2A). Overall, this signifies that the appendix contains an abundance of aggregation-prone alpha-synuclein proteoforms (*32–35*).

**Figure 2.**
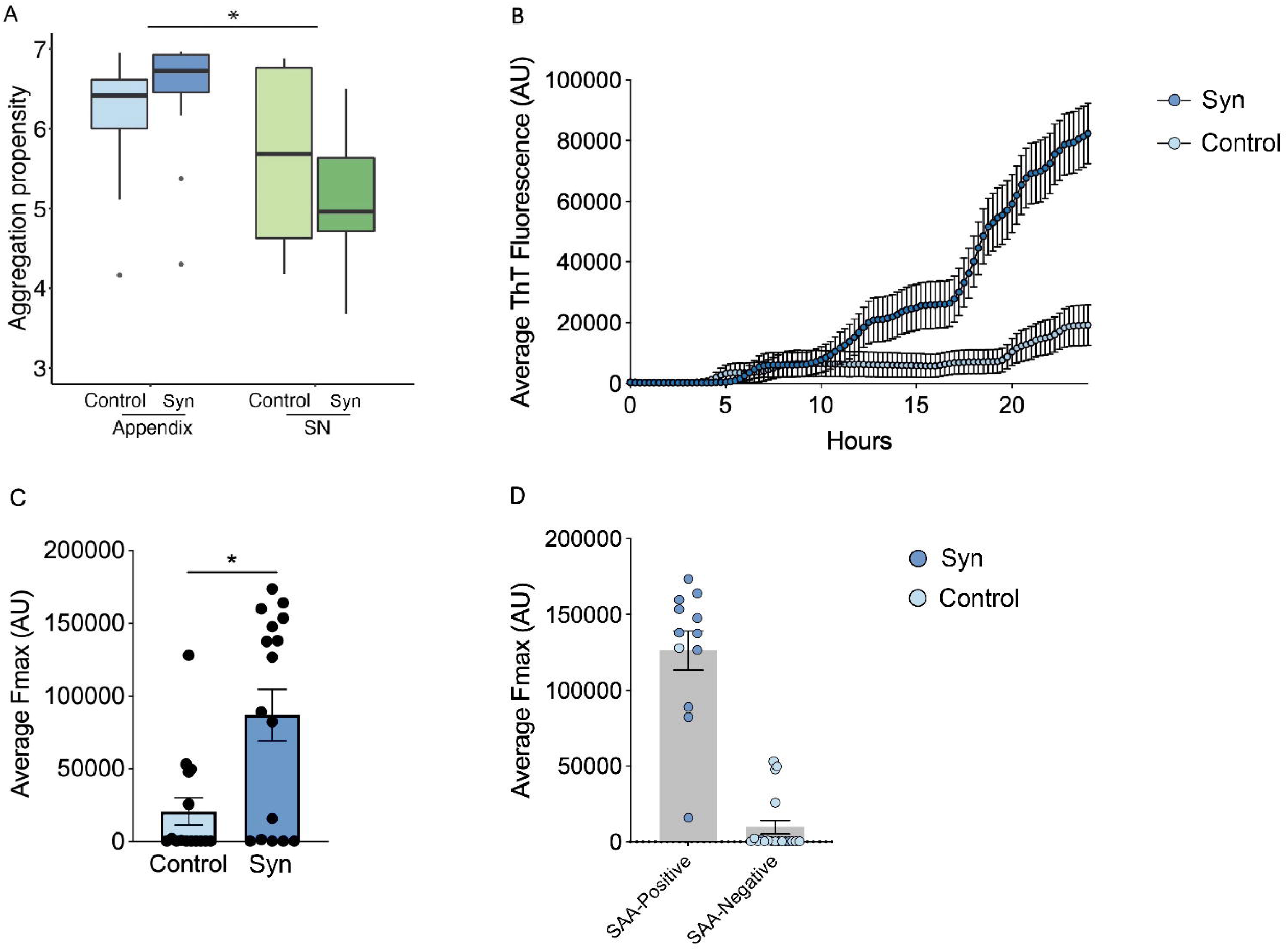
Aggregation propensity and seeding activity of alpha-synuclein. **(A)** High aggregation propensity of alpha-synuclein in the synucleinopathy (Syn) appendix compared to substantia nigra (SN). Aggregation propensity for each proteoform was estimated using PASTA 2.0. Robust linear regression was used for statistical analysis adjusting for diagnosis (synucleinopathy/control) and age, with aggregation propensity as dependent variable and tissue as independent variable. **(B)** Aggregation kinetics as measured by the assay demonstrated significantly increased average ThT fluorescence (AU) in synucleinopathy appendixes (n=16) over time compared to controls (n=15). This suggests the higher seeding activity of misfolded alpha-synuclein in synucleinopathy appendixes versus controls with no detectable seeding activity. **(C)** The maximum fluorescence (Fmax) recorded at the plateau of aggregation was significantly higher in synucleinopathy appendix as compared to controls (p<0.05). The graph includes data from all 3 replicates in each sample. Since positive/negative scores are defined by the number of replicates positive, this explains why four individuals have higher average fluorescence, but only one is considered positive. **(D)** In enhanced alpha-synuclein-SAA, positive seeding activity was detected in 11 synucleinopathy and 1 control appendix, while 5 synucleinopathy and 14 control appendices were negative for seeding activity. Mann-Whitney test was used to analyze average ThT fluorescence, Fmax SAA positive activity. Data are presented as mean ± SEM.

In parallel, we employed an enhanced alpha-synuclein-SAA to detect alpha-synuclein seeding activity in postmortem appendix tissues from synucleinopathy patients compared to matched controls. Using ThT fluorescence (AU) to measure seeding activity, we found 11 out of 16 (68.75%) Synucleinopathy appendices exhibited increased seeding activity (Figure 2D) as evidenced by increased ThT fluorescence (Figure 2B). In contrast, only 1 out of 15 controls (6.6%) demonstrated detectable seeding activity. Over a period of 24 h, synucleinopathy appendices showed a progressive increase in ThT fluorescence, whereas fluorescence in controls remained lower (Figure 2B). Furthermore, the average Fmax recorded at the plateau was significantly higher in synucleinopathy appendix as compared to controls (p=10^-4^) (Figure 2C). These results show significantly higher overall alpha-synuclein seeding activity in synucleinopathy appendix.

In the appendix and SN, we detected full-length alpha-synuclein lacking N-terminal acetylation, which has been shown to substantially increase its aggregation potential while reducing its lipid and synaptic vesicle binding affinity (*32–35*)(*p*<10^-4^; Figure 1D). Indeed, full-length alpha-synuclein that lacked N-terminal acetylation comprised 17.9% of the total full-length alpha-synuclein in the appendix and 3.8% in the SN. The only other N-terminal unacetylated alpha-synuclein proteoform detected in our study was unAc-alpha-synuclein^1-114^, which was primarily found in the appendix; present in 50.9% of appendix samples compared to only 21.1% of SN samples (unAc-alpha-synuclein^1-114^ present in 27 of 53 appendix samples and 4 of 19 SN samples; Figure 1D). Thus, delay in N-terminal acetylation in the appendix may accelerate alpha-synuclein aggregation in this tissue. No differences in N-terminal unacetylated alpha-synuclein were observed between synucleinopathy cases and controls (Supplementary Figure S4).

### Differential gene expression in disease

A total of 22029 genes were assessed, out of which 137 genes were significantly upregulated, and 113 genes were significantly downregulated in synucleinopathy appendix compared to healthy appendix (p-adj<0.05) (Figure 3A) (supplementary table S1). We identified dysregulation of genes related to heat shock and chaperone proteins associated with protein folding and degradation, immune response, as well as genes associated with cilia assembly and organization (Table 1).

**Figure 3.**
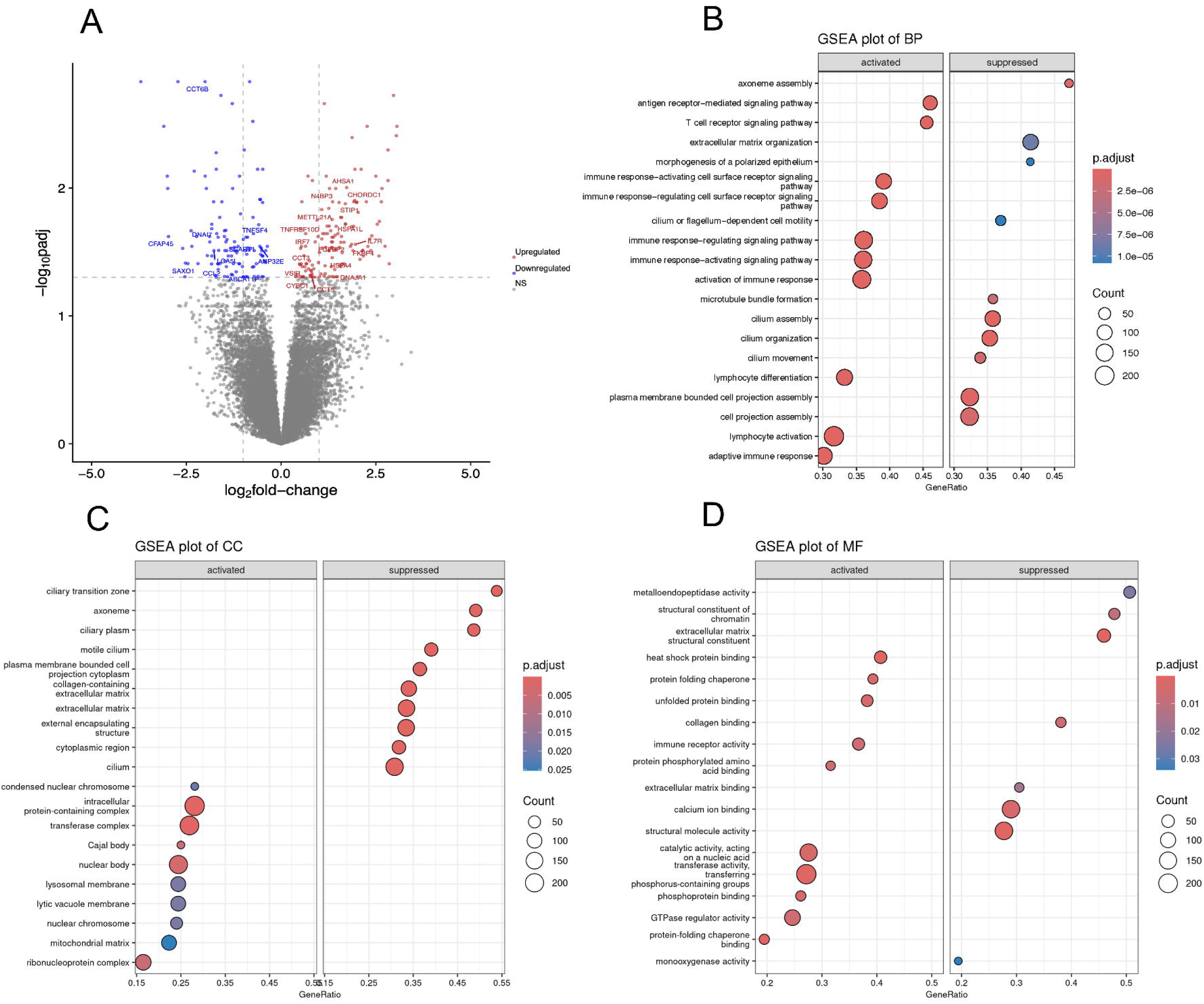
Dysregulation of genes and pathways in synucleinopathy patients’ appendices. **(A)** Volcano plot depicting differentially expressed genes in the appendix of synucleinopathy patients (n=11) as compared to controls (n=13). The upregulated (red) and downregulated (blue) genes with log_2_fold-change >2 (p-adj <0.05) are highlighted. The genes with no significant (NS) change are shown as grey dots. Gene Set Enrichment Analysis (GSEA) of Gene Ontology (GO) terms identified pathways affected by the DE genes in PD appendices vs controls. **(B)** GO-Biological Process, **(C)** GO-Cellular Components, and **(D)** GO-Molecular Function show activated (left) and suppressed (right) pathways in the PD appendix. Most of the DE genes and dysregulated pathways in synucleinopathy appendix contribute to immune responses, protein folding and cilia-related processes.

**Table 1.**
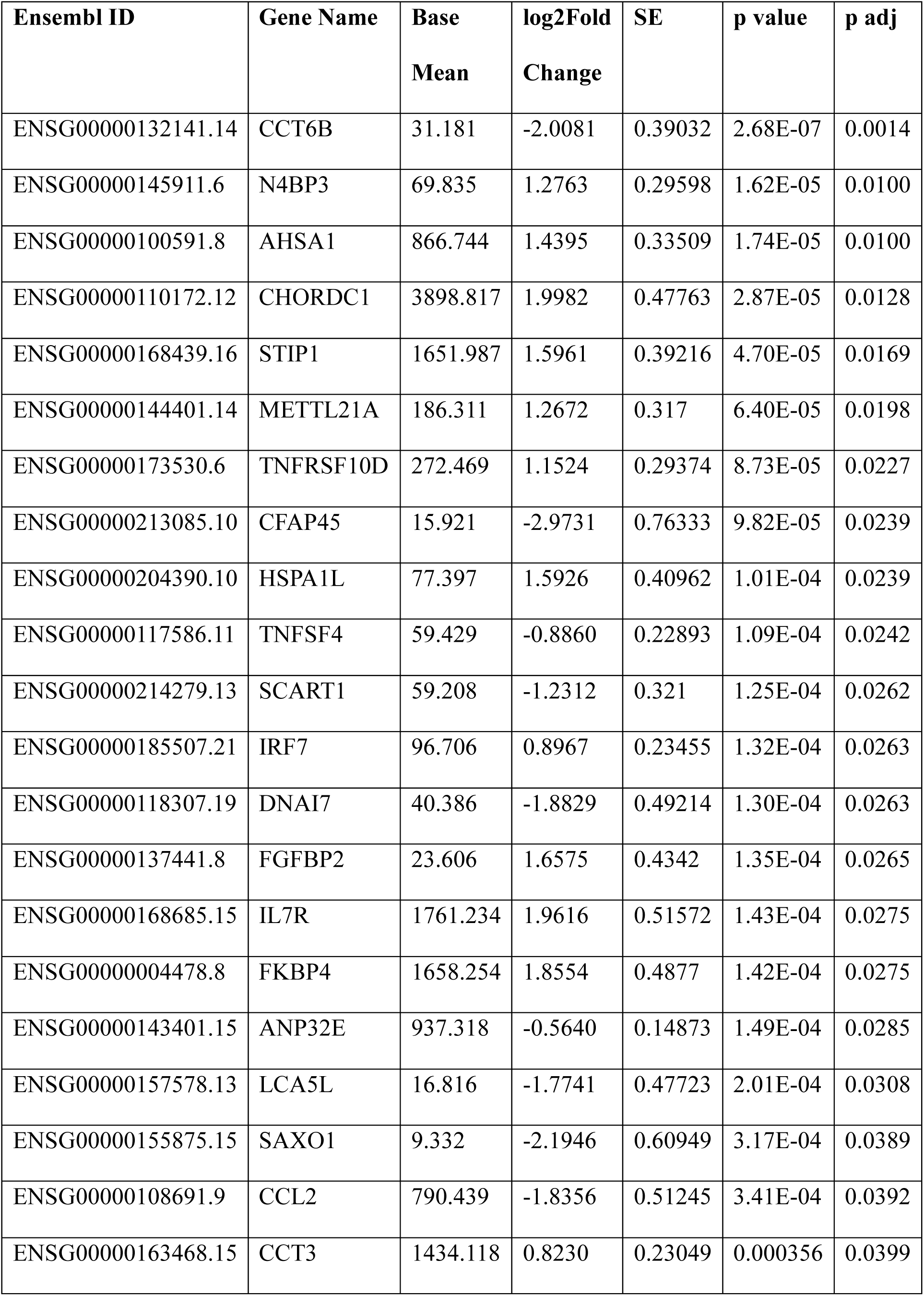
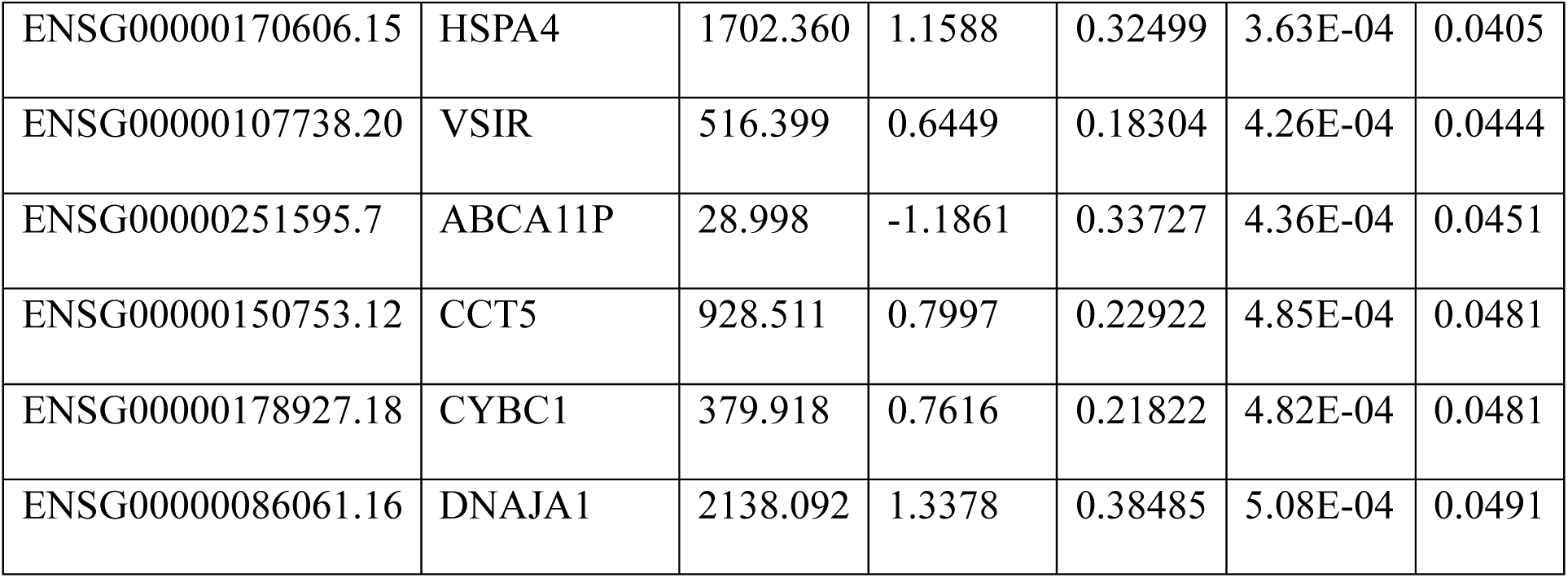
Subset of differentially expressed genes linked to synucleinopathy-associated processes in appendix samples from synucleinopathy patients vs. controls.

| **Ensembl ID** | **Gene Name** | **Base**<br>**Mean** | **log2Fold**<br>**Change** | **SE** | **p value** | **p adj** |
| --- | --- | --- | --- | --- | --- | --- |
| ENSG00000132141.14 | CCT6B | 31.181 | -2.0081 | 0.39032 | 2.68E-07 | 0.0014 |
| ENSG00000145911.6 | N4BP3 | 69.835 | 1.2763 | 0.29598 | 1.62E-05 | 0.0100 |
| ENSG00000100591.8 | AHSA1 | 866.744 | 1.4395 | 0.33509 | 1.74E-05 | 0.0100 |
| ENSG00000110172.12 | CHORDC1 | 3898.817 | 1.9982 | 0.47763 | 2.87E-05 | 0.0128 |
| ENSG00000168439.16 | STIP1 | 1651.987 | 1.5961 | 0.39216 | 4.70E-05 | 0.0169 |
| ENSG00000144401.14 | METTL21A | 186.311 | 1.2672 | 0.317 | 6.40E-05 | 0.0198 |
| ENSG00000173530.6 | TNFRSF10D | 272.469 | 1.1524 | 0.29374 | 8.73E-05 | 0.0227 |
| ENSG00000213085.10 | CFAP45 | 15.921 | -2.9731 | 0.76333 | 9.82E-05 | 0.0239 |
| ENSG00000204390.10 | HSPA1L | 77.397 | 1.5926 | 0.40962 | 1.01E-04 | 0.0239 |
| ENSG00000117586.11 | TNFSF4 | 59.429 | -0.8860 | 0.22893 | 1.09E-04 | 0.0242 |
| ENSG00000214279.13 | SCART1 | 59.208 | -1.2312 | 0.321 | 1.25E-04 | 0.0262 |
| ENSG00000185507.21 | IRF7 | 96.706 | 0.8967 | 0.23455 | 1.32E-04 | 0.0263 |
| ENSG00000118307.19 | DNAI7 | 40.386 | -1.8829 | 0.49214 | 1.30E-04 | 0.0263 |
| ENSG00000137441.8 | FGFBP2 | 23.606 | 1.6575 | 0.4342 | 1.35E-04 | 0.0265 |
| ENSG00000168685.15 | IL7R | 1761.234 | 1.9616 | 0.51572 | 1.43E-04 | 0.0275 |
| ENSG00000004478.8 | FKBP4 | 1658.254 | 1.8554 | 0.4877 | 1.42E-04 | 0.0275 |
| ENSG00000143401.15 | ANP32E | 937.318 | -0.5640 | 0.14873 | 1.49E-04 | 0.0285 |
| ENSG00000157578.13 | LCA5L | 16.816 | -1.7741 | 0.47723 | 2.01E-04 | 0.0308 |
| ENSG00000155875.15 | SAXO1 | 9.332 | -2.1946 | 0.60949 | 3.17E-04 | 0.0389 |
| ENSG00000108691.9 | CCL2 | 790.439 | -1.8356 | 0.51245 | 3.41E-04 | 0.0392 |
| ENSG00000163468.15 | CCT3 | 1434.118 | 0.8230 | 0.23049 | 0.000356 | 0.0399 |
| ENSG00000170606.15 | HSPA4 | 1702.360 | 1.1588 | 0.32499 | 3.63E-04 | 0.0405 |
| ENSG00000107738.20 | VSIR | 516.399 | 0.6449 | 0.18304 | 4.26E-04 | 0.0444 |
| ENSG00000251595.7 | ABCA11P | 28.998 | -1.1861 | 0.33727 | 4.36E-04 | 0.0451 |
| ENSG00000150753.12 | CCT5 | 928.511 | 0.7997 | 0.22922 | 4.85E-04 | 0.0481 |
| ENSG00000178927.18 | CYBC1 | 379.918 | 0.7616 | 0.21822 | 4.82E-04 | 0.0481 |
| ENSG00000086061.16 | DNAJA1 | 2138.092 | 1.3378 | 0.38485 | 5.08E-04 | 0.0491 |

Gene Set Enrichment Analysis (GSEA) was performed to assess whether predefined gene sets exhibit statistically significant differences between synucleinopathy and control samples. Differentially expressed (DE) genes in synucleinopathy appendix samples were found to be enriched in a total of 384 pathways, categorized as follows: 303 in GO Biological Processes (GO BP), 46 in GO Molecular Functions (GO MF), 31 in GO Cellular Components (GO CC), and 4 in KEGG pathways (supplementary data files S2-S5). The analysis highlighted several key processes involved in protein homeostasis and immune signaling that were significantly upregulated, and some related to ciliary dynamics that were downregulated (Figure 3B, 3C, 3D).

### Differential DNA methylation in disease

The overall distribution of differentially methylated regions (DMRs) across gene regions, CpG regions, and the total methylation profile suggests a predominantly hypomethylated status in synucleinopathy appendix samples compared to healthy controls (Figure 4B, 4C, 4D). Most CpG sites were at promoter region transcription start site 1500 (TSS1500)(67.29%), followed by 5’ untranslated region (5’UTR)(12.87%), transcription start site 200 (TSS200)(10.99%), gene body (6.43%), intergenic (1.61%) and 1^st^ Exon (0.80%). In terms of CpG regions, the highest number of CpG sites was at islands (68.90%), followed by shores (22.52%), open sea (6.97%), and shelves (1.61%).

**Figure 4.**
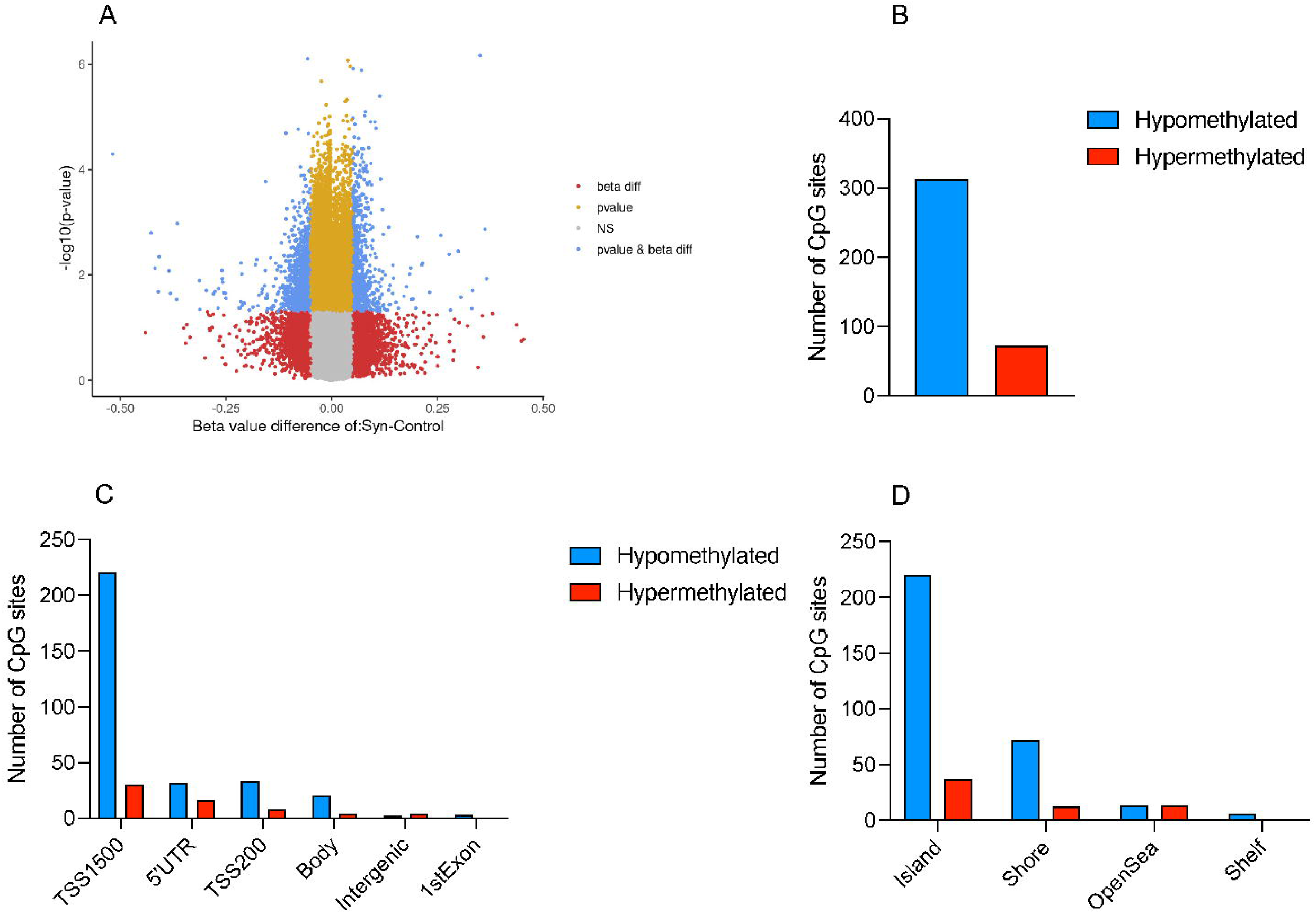
Epigenetic dysregulations in synucleinopathy patients’ appendices. **(A)** The volcano plot displays differential methylation between synucleinopathy appendices (n=11) and controls, (n=13) with the x-axis showing beta value differences (synucleinopathy – control). Blue dots represent observations that were significantly different in both p-value and beta value. **(B)** The total number of hypomethylated and hypermethylated CpG sites exhibiting more hypomethylation in synucleinopathy appendix as compared to controls. **(C)** The distribution of number of CpG sites across various gene regions in synucleinopathy appendices compared to controls. **(D)** The distribution of number of CpG sites across CpG regions in synucleinopathy appendices compared to controls.

We found 46 unique genes in the vicinity of 45 DMRs-39 genes were hypomethylated, while 7 genes were hypermethylated (supplementary table S6). Methylation differences between synucleinopathy and control appendices were subtle but aligned with our transcriptomic analysis; we found differentially methylated genes related to protein degradation, folding and homeostasis, immune response, and ciliary and cytoskeletal dynamics (Table 2). Additionally, we found differential methylation in some genes associated with oxidative stress and mitochondrial function, both of which have been implicated in neurodegeneration. GSEA revealed neuronal signaling and development pathways that were enriched for hypomethylated genes, and stress response and cell signaling pathways that were enriched for hypermethylated genes in synucleinopathy appendix (Figure 5) (supplementary tables S7-S10).

**Table 2.** Subset of differentially methylated genes linked to synucleinopathy-associated processes in appendix samples from synucleinopathy patients vs. controls.

| tpCombine | Seg Chrm | Seg Estimate | p value | p adj | Relation to<br>Gene<br>Category |
| --- | --- | --- | --- | --- | --- |
| MIP | chr12 | -0.056460768 | 7.8464E-07 | 0.02640 | 5'UTR |
| H19 | chr11 | -0.045129478 | 5.1411E-12 | 2.2613E-06 | TSS1500 |
| GLI4 | chr8 | -0.040098331 | 1.5587E-06 | 0.02981 | TSS1500,<br>Body |
| GAS7 | chr17 | -0.031342502 | 2.5995E-06 | 0.0420 | TSS1500,<br>5'UTR |
| CDK16 | chrX | -0.030810321 | 1.1623E-09 | 2.5563E-04 | TSS1500,<br>TSS200 |
| TBL1X | chrX | -0.029706597 | 2.7826E-08 | 0.0041 | TSS1500,<br>TSS200,<br>Body |
| TMC4 | chr19 | -0.02341605 | 2.9657E-06 | 0.0432 | TSS200 |
| EIF1AX | chrX | -0.011564424 | 1.4196E-06 | 0.0284 | 5'UTR |
| WBP11 | chr12 | -0.00582025 | 5.0605E-06 | 0.0495 | TSS1500 |
| NR4A1 | chr12 | -0.005543784 | 4.9188E-06 | 0.0495 | Body |
| MRPL41 | chr9 | -0.005405291 | 4.7806E-07 | 0.0264 | 1stExon,<br>Body |
| TEX22 | chr14 | -0.005314138 | 3.9308E-07 | 0.0264 | TSS1500 |
| MTA1 | chr14 | -0.005314138 | 3.9308E-07 | 0.0264 | TSS1500 |
| SH3GLB1 | chr1 | -0.005202031 | 2.0330E-06 | 0.0358 | TSS1500,<br>Body |
| AKR7A2 | chr1 | -0.005023715 | 4.0374E-06 | 0.0487 | TSS1500 |
| SLC66A1 | chr1 | -0.005023715 | 4.0374E-06 | 0.0487 | TSS1500 |
| ANXA5 | chr4 | -0.004686528 | 3.4229E-06 | 0.0470 | TSS200,<br>5'UTR |
| ZMYND11 | chr10 | -0.004486106 | 6.0076E-07 | 0.0264 | TSS1500,<br>TSS200,<br>Body |
| DPH3 | chr3 | -0.004301509 | 1.1084E-06 | 0.0284 | TSS1500 |
| OXNAD1 | chr3 | -0.004301509 | 1.1084E-06 | 0.0284 | TSS1500 |
| UBTF | chr17 | -0.003862467 | 2.7495E-06 | 0.0420 | TSS1500 |
| NAP1L1 | chr12 | -0.003581127 | 8.3028E-07 | 0.0264 | TSS1500,<br>TSS200,<br>5'UTR |
| ACBD3 | chr1 | -0.003069002 | 3.9148E-06 | 0.0487 | 5'UTR |
| ABCB9 | chr12 | -0.003039661 | 8.9398E-08 | 0.0079 | TSS1500 |
| LSM2 | chr6 | -0.002908889 | 4.2098E-06 | 0.0487 | TSS1500,<br>TSS200,<br>5'UTR |
| DHX29 | chr5 | -0.002533829 | 1.0731E-06 | 0.0284 | TSS1500 |
| MTREX | chr5 | -0.002533829 | 1.0731E-06 | 0.0284 | TSS1500 |
| TRIM33 | chr1 | -0.002213311 | 1.3577E-06 | 0.0284 | 5'UTR |
| TOR3A | chr1 | -0.00137959 | 3.8652E-06 | 0.0487 | TSS200,<br>Body |
| DNMT3A | chr2 | -1.27370184049748E-04 | 4.1074E-06 | 0.0487 | TSS1500,<br>5'UTR |
| TRNP1 | chr1 | -5.70366549201819E-05 | 4.7327E-06 | 0.0495 | TSS1500,<br>TSS200, 1st<br>Exon, Body |
| PPP2R2B | chr5 | 0.019478798 | 2.3741E-06 | 0.0402 | TSS1500,<br>TSS200,<br>5'UTR |
| R3HDM2 | chr12 | 0.033485722 | 5.0267E-06 | 0.0495 | 5'UTR |
| DCUN1D1 | chr3 | 0.071012065 | 1.2839E-06 | 0.0284 | 5'UTR |
| MEI1 | chr22 | 0.351120112 | 6.7254E-07 | 0.0264 | Body |

**Figure 5.**
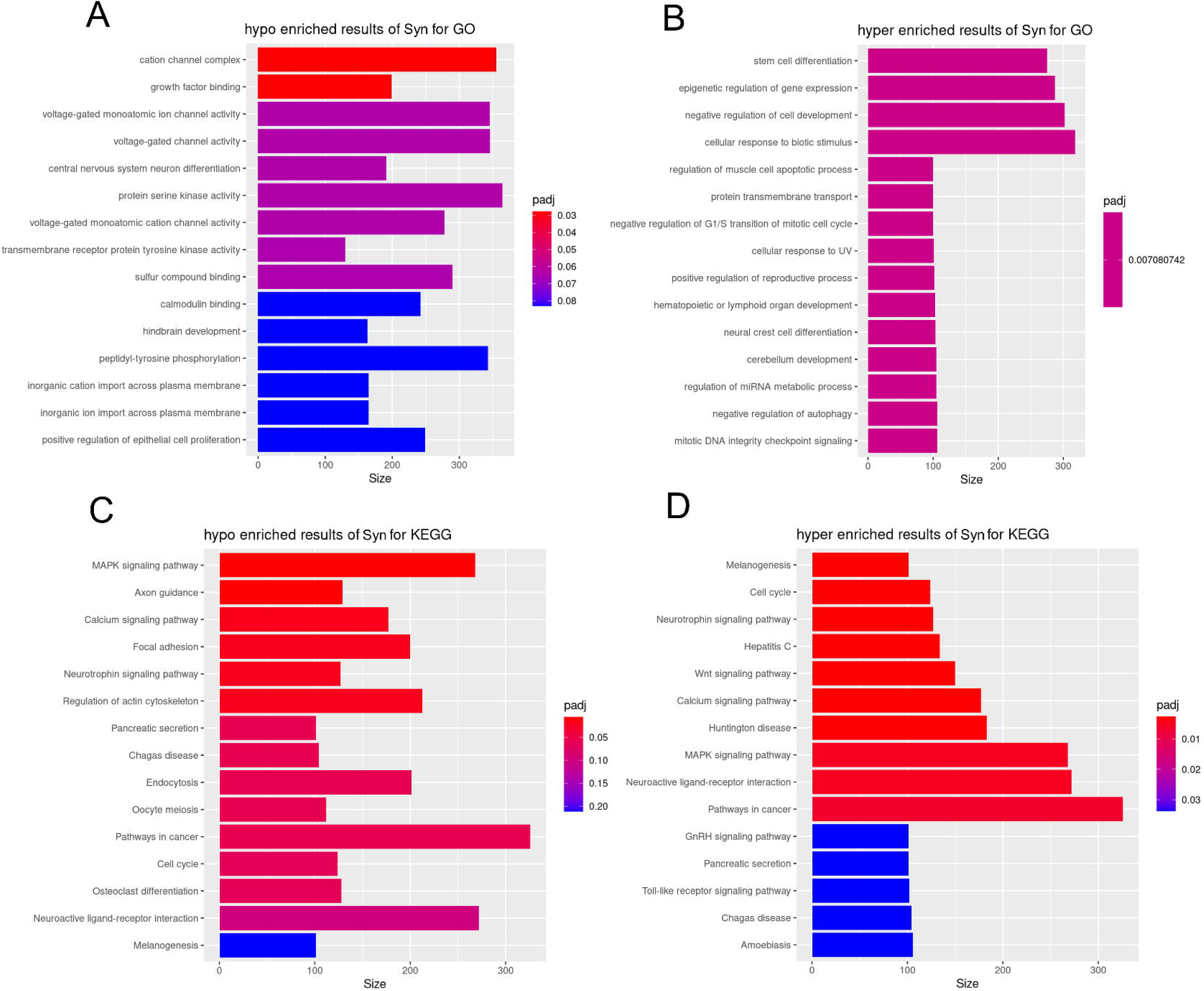
Gene Set Enrichment Analysis (GSEA) of Gene Ontology (GO) terms identified pathways affected by the DM genes in the synucleinopathy appendices vs controls. **(A)** Hypomethylated pathways mapped to GO database, **(B)** hypermethylated pathways mapped to GO database. **(C)** Hypomethylated pathways mapped to KEGG database, **(D)** hypermethylated pathways mapped to KEGG database.

### Functional clustering based on protein interactions in disease

Differentially expressed and methylated genes (p-adj<0.05) from RNA-seq and DNA methylation analyses were analyzed using the STRING database to create a network map (Figure 6). Functional clusters identified through Markov Cluster Algorithm (MCL) aligned with key pathways highlighted in GSEA analysis, including protein homeostasis (Unfolded protein binding and Chaperone complex), immune response (Autoimmune disease), and ciliary structure (Axonemal microtubule) networks.

**Figure 6.**
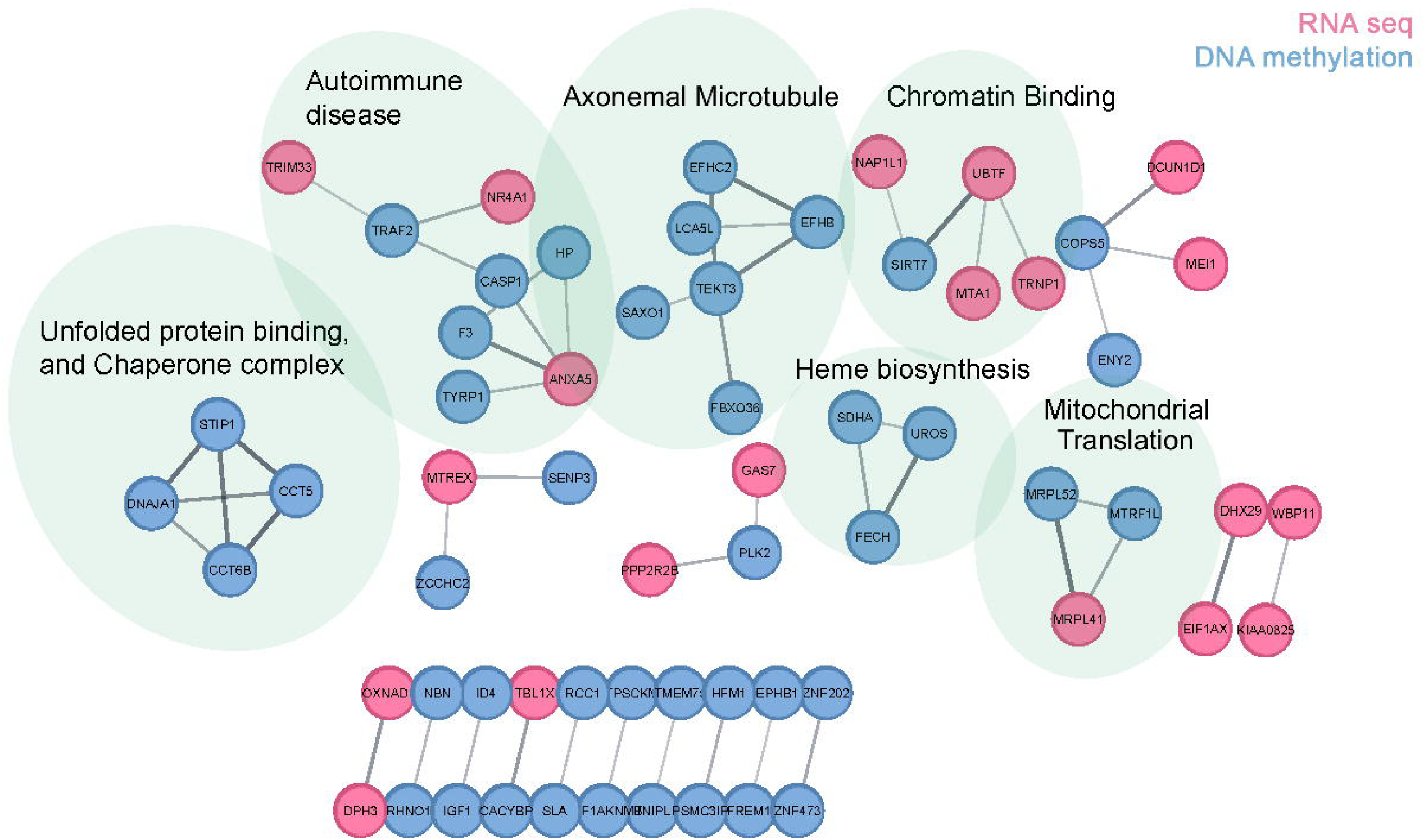
Protein–protein interaction network of significantly altered genes in synucleinopathy appendix. Significant genes (p-adj. 0.05) identified from RNA-Seq and DNA methylation analysis were analyzed using the STRING database. The plot excludes genes that lack STRING-annotated functional connectivity. Nodes are colored according to whether the gene was identified via RNA or DNA methylation analysis. Markov Cluster Algorithm (MCL) was used to organize nodes, and the top statistically significant functional enrichment (i.e., lowest FDR value) was displayed above each cluster.

## DISCUSSION

The vermiform appendix has emerged as a potential peripheral site where increased levels of aggregated alpha-synuclein potentially appear early in the course of synucleinopathies (*9*). In the presence of impaired autophagy-lysosomal function in the same tissue (*13*), aggregated alpha-synuclein might propagate from enteric nerves to the CNS, making the appendix one possible site for initiation of the first steps in the pathogenesis of synucleinopathies. Despite growing evidence, the mechanisms causing alpha-synuclein aggregation in the appendix are poorly understood. To address this knowledge gap, we employed proteomic, transcriptomic, and epigenomic profiling of appendices from synucleinopathy patients and healthy controls. Our findings reveal a distinct, aggregation-prone pool of alpha-synuclein proteoforms in the synucleinopathy appendix, and significant dysregulation in protein homeostasis, immune response, and ciliary function and processes in the synucleinopathy appendix. These findings provide a novel molecular insight into the vermiform appendix as a peripheral site where alpha-synuclein aggregates in the earliest stages of the disease process.

To characterize the molecular features underlying synuclein pathology, we investigated alpha-synuclein aggregation and truncation in appendix tissues and SN. When we modeled aggregation propensity for the total pool of alpha-synuclein proteoforms identified using TD-MS, we observed a higher aggregation propensity in the appendix in comparison to SN. This suggests the unique proteolytic processing in the vermiform appendix “primes” the endogenous alpha-synuclein pool for pathological aggregation, which may also be true of other peripheral tissues where pathology has also been documented (*36, 37*). We then investigated this experimentally using alpha-synuclein-SAA, where we detected seeding activity in 68.75% of synucleinopathy appendix samples compared to 6.6% of healthy appendix samples. Importantly, fluorescence kinetics revealed higher seeding activity, reflected by greater ThT signal and Fmax, in synucleinopathy appendix relative to controls. These findings are consistent with previous studies showing that gastrointestinal tissues from synucleinopathy patients exhibit more consistent seeding activity compared to controls (*38*). Notably, our finding that 68.75% of synucleinopathy appendices are SAA-positive aligns well with the proposed “body-first subtype” of Lewy body diseases (*39*), in which pathology is hypothesized to begin in the enteric nervous system and ascend to the brain via the vagus and sympathetic nerves. According to this model, approximately half of the Lewy body disease patients belong to the “body-first” subgroup, and the remaining fall into a “brain-first” category, with around 70% cases of dementia with Lewy bodies (DLB) and only ∼30% of PD cases exhibiting a body-first pattern (*40*). Since our synucleinopathy cohort was DLB-rich (n=12 of 16), the observation that 68.75% of our synucleinopathy appendix samples were SAA-positive is consistent with the expected proportion of body-first cases. Conversely, 31.25% of synucleinopathy appendices were negative for seeding activity in alpha-synuclein-SAA, suggesting a brain-first trajectory while exhibiting less peripheral involvement, demonstrating the heterogeneity of alpha-synuclein pathology within synucleinopathies.

We previously identified the appendix as a reservoir of intraneuronal alpha-synuclein aggregates that are enriched with truncated proteoforms (*9*), which raised questions about the proteolytic turnover of alpha-synuclein in the appendix. Here we addressed those questions by cataloging all detectable alpha-synuclein proteoforms using a TD-MS approach, similar to what has been done in brain tissue (*26*). This study provides one of the most comprehensive catalogs of alpha-synuclein truncation in human tissues to date. We compared alpha-synuclein proteoforms between tissues and found proteoform abundance varied significantly between the appendix and SN. Unexpectedly, 9 proteoforms were unique to the appendix and were not detected in any of the SN samples. Conversely, two proteoforms, namely alpha-synuclein^78-113^ and alpha-synuclein^ac1-122^, were decreased in the synucleinopathy appendix compared to controls. The decrease in these proteoforms can be attributed to either abnormal proteolytic turnover of alpha-synuclein in the synucleinopathy appendix or possibly indicative of aggregation, as proteoforms that form insoluble inclusions would not be detected with our methodology.

Several of the cleavage sites and proteoforms we identified have been described previously (*9, 21, 26*), but we also characterized proteoforms not yet described in literature. As truncated forms of alpha-synuclein are a substantial component of inclusions (*41*), characterization of the proteoforms present in the appendix and SN may improve our understanding of the origins of synucleinopathies. Truncation and N-terminal acetylation were the only detectable PTMs. Like many eukaryotic proteins, the N-terminal methionine of mature alpha-synuclein is permanently acetylated. Here, we made the interesting observation that full-length and some C-terminally truncated alpha-synuclein species were unacetylated. Although the levels of N-terminally unacetylated alpha-synuclein were relatively low in the brain and appendix, several appendix samples showed a surprising abundance of this particular proteoform (∼1:2). Furthermore, unacetylated alpha-synuclein was the third most abundant proteoform we detected regardless of synucleinopathy or tissue type. Unacetylated alpha-synuclein likely represents immature or newly synthesized protein that has not yet been enzymatically modified. Thus, in some instances alpha-synuclein was cleaved prior to becoming a mature protein. This raises the possibility that alpha-synuclein truncation can be an early event in the protein’s natural lifecycle. Of note, we did not detect any phosphorylated proteoforms, including the disease-associated phosphoserine 129, which is in agreement with previous studies showing very low levels of this modification in the normal soluble alpha-synuclein pool (*26*). Furthermore, alpha-synuclein from the human appendix was more abundant in truncated proteoforms when compared to the brain. Combined with our previous observation of an impaired lysosomal autophagy system in the Parkinson’s disease appendix (*13*), this might suggest that the turnover of alpha-synuclein was impaired, leading to an accumulation of truncated proteoforms and the formation of proteopathic seeds that could initiate the first steps of synucleinopathy.

Together, our results suggest that proteolytic degradation (i.e. turnover) of alpha-synuclein is tissue-specific, and therefore some tissues, like the vermiform appendix, may be more likely to accumulate alpha-synuclein, including rare truncated proteoforms. Some important considerations should be taken when interpreting our findings. First, immunopurification of alpha-synuclein was conducted using clone 42 anti-alpha-synuclein antibody (BD Transduction Laboratories) which maps to residues 91-99 of human alpha-synuclein (*42*), and thus, all proteoforms detected expectedly contain this epitope, and any proteoforms lacking this epitope would not be detected here. The total alpha-synuclein signal intensity was similar between appendix and SN samples despite relatively low expression of alpha-synuclein in the appendix, indicating that immunopurification was conducted at near-saturating conditions. As a result, our data are not indicative of changes in total alpha-synuclein content of the tissues but instead are an indicator of the proteoform signature of the samples. Because our analyses were performed on the triton x100 soluble fraction of alpha-synuclein, it is limited to proteoforms found outside of inclusion bodies. Conceptually, proteoforms identified here are “pre-pathological” and provide insight into the normal turnover of this protein in peripheral and CNS tissues.

Building on our TD-MS and seed amplification assay results, which indicate a heightened aggregation potential in synucleinopathy appendix, we turned to multi-omics profiling to investigate the transcriptional and epigenetic underpinnings of this phenomenon. We identified a prominent alteration in genes involved in proteostasis and quality control, which suggests cellular stress and activated mechanisms to counteract misfolded protein burden in the synucleinopathy appendix. Activated pathways associated with molecular assemblies such as intracellular protein-containing complex, transferase complex and ribonucleoprotein complex hint towards changes in protein turnover, potentially instigated by aggregation of misfolded alpha-synuclein in the synucleinopathy appendix. Molecular function pathways relevant to synuclein pathology such as protein phosphorylated amino acid binding, phosphoprotein binding, and phosphotransferase activity were also upregulated. In the context of PD, these pathways are central to the recognition of phosphorylated alpha-synuclein, and binding to 14-3-3 domain proteins to produce Lewy body pathology (*43*), further highlighting a dysregulated proteostasis network. Simultaneously, protein folding chaperone, and HSP/protein folding chaperone binding pathways were upregulated in synucleinopathy appendix, suggesting a cellular attempt to attenuate misfolding and aggregation of alpha-synuclein. Activation of pathways associated with lysosomal membrane and lytic vacuole membrane indicates an increase in compensatory autophagic clearance. DNA methylation pathway enrichment revealed hypomethylated (may indicate potentially increased activity) genes associated with cation channel complex, as well as differential methylation of calcium signaling pathway genes (both hypo- and hyper-enriched), reflecting altered regulation of processes implicated in alpha-synuclein aggregation (*44*). Additionally, hypermethylation (may indicate potentially decreased activity) of genes associated with Wnt signaling pathway and negative regulation of autophagy is consistent with our previous observations at transcriptomic level. At the gene level, these changes were reflected in the observed upregulation of several molecular chaperones and co-chaperones of heat shock proteins (HSPs) such as AHA1, STIP1, METTL21A, HSPA1L, and HSPA4 (*45*). The chaperones HSP70 and HSP90 play a crucial role in mitigating cellular stress such as accumulation of misfolded proteins (*46*), degradation of toxic protein aggregates (*47*), and modulating alpha-synuclein dynamics (*48*). We also observed hypomethylation in promoter regions of several genes implicated in autophagy, lysosomal trafficking, and stress signaling, such as CDK16, SH3GLB1, and ABCB9 in the synucleinopathy appendix (*49*) (*50*) (*51*), and hypermethylation of DCUN1D1, which was previously identified as a novel candidate gene in protein interactome-based PD studies (*52*).

Immune response-related pathways such as T cell receptor signaling, lymphocyte differentiation and activation, and adaptive immune response, were activated in transcriptomic enrichment analysis, indicating a sustained state of immune activation in the synucleinopathy appendix. Interestingly, immune response regulating pathways were also activated, potentially suggesting a compensatory attempt to maintain immune balance in response to proteostatic stress. Differential methylation of the stress and immune regulatory MAPK signaling pathway (both hypo- and hyper-enriched), and hyper-enrichment of Toll-like receptor signaling also suggest immune remodeling at the epigenetic level. Our observations at gene level, such as upregulation of chemokine and interferon response genes CCL2, IRF7, IL7R, and the immunomodulatory checkpoint gene VSIR, further reflect the complex immune environment within synucleinopathy appendixes. Particularly, in the context of neurodegenerative disease, CCL2 and VSIR have been associated with microglial activation and neuroinflammation (*53, 54*). We also found hypomethylation in the promoter region of long noncoding RNA H19, the overexpression of which has been associated with downregulation of tight junction proteins ZO-1 and occludin (*55*). This may, in turn, compromise the integrity of the epithelial barrier in the synucleinopathy appendix and further increase vulnerability to inflammation, while simultaneously allowing for microbial products inside the appendix to circulate systemically. This so-called ‘leaky gut’ state, in combination with the perturbed local immune environment, may contribute to systemic inflammation and potentially influence alpha-synuclein pathology (*56*). These results are also consistent with another recent study, where transcriptomic and epigenomic profiling of the colonic mucosa revealed shared immune dysregulation in PD and inflammatory bowel disease patients (*57*). Their findings also suggest that chronic peripheral inflammation can increase the risk for PD in inflammatory bowel disease patients, further strengthening the link between gut immune dysregulation and neurodegeneration.

Our data also revealed consistent dysregulation of genes related to primary and motile ciliary structure and function. Primary cilia are essential cellular organelles that are present on the epithelial layer, as well as in the enteric neurons embedded inside the appendix (*58*). These structures are involved in modulating the Sonic Hedgehog (Shh) pathway, which is crucial for neurogenesis, and has been linked to PD (*59, 60*). Motile cilia, on the other hand, are found on specialized cells and facilitate the movement of fluid over epithelial surfaces (*61*). In pathway analysis, we observed significant disruption of cilia-related processes as well as other structural components, as reflected by suppressed pathways associated with axoneme assembly, cilium assembly and organization, ciliary plasm, mobile cilium, ciliary transition zone, cilium- or flagellum-dependent cell motility, epithelial morphogenesis, and extracellular matrix. Beyond dysfunctional signaling, disruption of these components would also have implications for barrier integrity, corroborating our earlier observation. The hypo-enrichment of KEGG terms such as regulation of actin cytoskeleton and focal adhesion may indicate compensatory activity of the epigenetic machinery, which appears to mitigate transcriptomic suppression. The observed downregulation of primary cilia stability gene SAXO1, and motile cilia function genes LCA5L, CFAP45, and DNAI7, as well as hypomethylation of the downstream cellular regulators GLI4 and GAS7, would further hint towards downstream disruption of Shh signaling and possibly, GI motility, in synucleinopathy patients. Interestingly, GLI4 and GAS7 have been associated with PD in previous studies (*62, 63*). Additionally, LRRK2 mutations, which are common in PD, have also been found to interfere with cilia formation and Shh signaling, further suggesting a potential mechanistic link between ciliary dysfunction and PD development (*64*).

Although we observed no direct overlap between differentially methylated genes and differentially expressed genes in our data after FDR correction, both datasets overlap substantially in functional themes. This suggests that even subtle methylation changes in the synucleinopathy appendix observed here could potentially influence gene expression, and the divergence could be because of additional regulatory layers such as histone modifications or non-coding RNAs.

The present study provides new insights into the molecular landscape of the vermiform appendix in PD, highlighting it as a potential contributor to pathology. The interplay between impaired protein homeostasis, immune dysregulation, and ciliary dysfunction may prove conducive to aggregation of alpha-synuclein, which could then lead to propagation of pathology to the central nervous system. Further research into the identified dysregulated genes and pathways, as well as aberrant alpha-synuclein processing in the appendix, may provide critical insights and inform the development of therapeutics for synucleinopathies.

## MATERIALS AND METHODS

### Study design

This study was designed to investigate the molecular mechanism underlying alpha-synuclein aggregation in vermiform appendix. We conducted TD-MS to characterize alpha-synuclein proteoforms in post-surgical appendix (24 synucleinopathy cases and 31 controls) and postmortem SN (10 synucleinopathy cases and 10 controls) tissue samples (20-100 mg). Next, we performed *in silico* modeling of aggregation propensity for each proteoform characterized in the previous experiment. We also investigated this *in vitro* utilizing alpha-synuclein-SAA to differentiate seeding activity exhibited by synucleinopathy and healthy control appendixes (synucleinopathy=16, control=15, 30-40mg each). Excluding the samples without genomic profiling permissions, the same appendix samples were utilized to explore gene regulation abnormalities through RNA seq and DNA methylation profiling (synucleinopathy=11, control=15, 30-40mg each). After initial QC, one healthy control appendix sample was excluded from subsequent analyses (final control n=14). Network analysis was performed to integrate findings from transcriptomic and epigenomic analyses. Due to sample collection constraints, no statistical methods were used to predetermine sample size. Number of technical replicates and positivity criteria for alpha-synuclein-SAA experiment are detailed below.

### Appendix sample processing for enhanced alpha-synuclein seed amplification assay (alpha-synuclein-SAA)

Postmortem human vermiform appendix samples were cryopulverized using the CP02 Automated Dry Pulverizer (Covaris LLC, Woburn, MA, USA) and prepared at a 3% (w/v) concentration using a homogenization buffer containing PBS and 1x EDTA-free protease inhibitor from Roche. Homogenization was performed with Precellys Lysing Kit using the Bertin Precellys-24 Dual homogenizer for two 30-second cycles at 6500 rpm. Following homogenization, the appendix samples were centrifuged at 800 × g for 1 minute at 4°C. The resulting supernatant was collected, vortexed, aliquoted, snap-frozen, and stored at -80°C until further use.

### Alpha-synuclein-SAA procedure

The enhanced alpha-synuclein-SAA used here is a modification of a previously described assay (*65*), and was performed following a protocol described by Ma et al. (*66*). Briefly, the reaction mixture contained 100mM PIPES pH 6.50, 500mM NaCl, 10µM thioflavin T (ThT), 0.1% sarkosyl, and 0.3mg/mL recombinant alpha-synuclein. The assay was performed in 96-well clear-bottom plates in the presence of two 3.2mm Si3N4 beads per well. Plates were incubated at 42°C with intermittent orbital shaking (800rpm) for 1 min every 15 mins and fluorescence was read at every cycle (15 min), using FLUOstar Omega (BMG LABTECH, Ortenberg, Germany). Samples were tested in triplicate during a 24h period and were considered positive when three replicates exhibited a fluorescence >1000 AU. Samples with 0 or 1 positive replicates were designated as negative. Samples with 2 positive replicates were considered inconclusive and were tested again. The maximum fluorescence (Fmax) of each sample was determined by averaging Fmax from all three replicates.

### Alpha-synuclein immunoprecipitation for mass spectrometry

Appendix and SN tissue samples were ground on liquid nitrogen with a mortar and pestle. The resulting powdered tissue was suspended in lysis buffer (PBS, 1% Triton X-100, 2mM EDTA, 2mM PMSF, 1X Roche protease inhibitor cocktail) and sonicated at 50% intensity with 2s pulses for 30 pulses total, on ice using Q55 model sonicator (QSonica). Samples were then vortexed and then incubated on ice for 30 min. Samples were centrifuged at 22,000-x *g* for 20 min, and the supernatant was retained. Protein content was determined by BCA assay (Thermo Fisher) and adjusted to 2 mg protein / mL lysis buffer.

For each sample, 50 µl of the tosylactivated beads (see supplementary methods for details on preparation of beads) was collected on a magnetic stand and combined with 200 µl of tissue lysate. Samples were then incubated overnight at 4°C with nutation. The sample-exposed beads were magnetically collected and washed three times with ice-cold lysis buffer (1 mL of PBS, 1% Triton X-100, 2mM EDTA, 2mM PMSF, 1 X Roche protease inhibitor cocktail), with each wash performed at 4°C with nutation for 30 min. Next, the samples were washed with 1 mL PBS pH 7.4, three times at 4°C with nutation for 15 min and then rinsed three times with 1 mL ultrapure water (Thermo Fisher). To elute alpha-synuclein from the beads, samples were suspended in 10 µL of 1% formic acid in ultrapure water and mixed vigorously for 5 min at room temperature. Samples were then centrifuged for 5 s, placed on the magnet, and the eluent collected into a protein LoBind Eppendorf tube (Sigma-Aldrich). The elution procedure was repeated with another 10 µL of 1% formic acid, and both eluents were pooled. The presence of alpha-synuclein was verified in an aliquot (2 µL) of the eluent by western blotting, and the remaining eluent was immediately snap frozen in liquid nitrogen and stored at -80°C until top-down mass spectrometry analysis.

### Liquid chromatography-mass spectrometry

After immunoprecipitation, samples eluted in high acid were loaded in a nanocapillary liquid chromatography (LC) (Dionex RSLCnano) setup for proteoform separation. A trap column (2 cm × 150 μm i.d.) and analytical (20 cm × 75 μm) column were used, both packed with PLRP-S stationary phase (5 μm particle size, Agilent, Santa Clara, CA) and proteins were eluted over a 40 min gradient of 95:5 water/acetonitrile with 0.2% formic acid to 5:95 water/acetonitrile into a custom electrospray ionization (ESI) source terminated with a PicoTip spray emitter (New Objective, Woburn, MA). Proteoforms were then analyzed online on a LTQ Velos Orbitrap Elite mass spectrometer (Thermo Fisher Scientific) operated in a data-dependent mode, using established instrument methods (*67*). Two LC-MS replicates of each sample were analyzed by LC-MS, and sample replicates were run in full random order to mitigate the effect of instrumental drift in differential proteoform quantification. The ensuing raw data was then searched for target alpha-synuclein proteoforms using a previously described custom MS1-based isotopic fitting software (see supplementary methods for details on data processing) (*68*).

### RNA sequencing (RNA-seq)

Total RNA was extracted from previously mentioned appendix samples utilized for alpha-synuclein-SAA using the KAPA RNA HyperPrep Kit (Kapa Biosystems, Wilmington, MA USA). Libraries were prepared by the Van Andel Genomics Core from 445 ng of the total RNA. Ribosomal RNA material was reduced using the QIAseq FastSelect –rRNA HMR Kit (Qiagen, Germantown, MD, USA). RNA was sheared to 300-400 bp. Prior to PCR amplification, cDNA fragments were ligated to IDT for Illumina TruSeq UD Indexed adapters (Illumina Inc, San Diego CA, USA). Quality and quantity of the finished libraries were assessed using a combination of Agilent DNA High Sensitivity chip (Agilent Technologies, Inc.), QuantiFluor® dsDNA System (Promega Corp., Madison, WI, USA), and Kapa Illumina Library Quantification qPCR assays (Kapa Biosystems). Individually indexed libraries were pooled and 100 bp paired-end sequencing was performed on an Illumina NovaSeq6000 sequencer to an average depth of 60M raw paired-reads per transcriptome. Base calling was done by Illumina RTA3 and output of NCS was demultiplexed and converted to FastQ format with Illumina Bcl2fastq v2.20.0. Differential gene expression (DGE) analysis was performed using DESeq2 (*69*) based on gene counts generated from STAR. Further details on data processing have been provided in supplementary methods.

### DNA methylation analysis

DNA extracted from previously mentioned cryopulverized appendix samples was quantified by Qubit fluorometry (Life Technologies), and 250 ng of DNA from each sample was bisulfite converted using the Zymo EZ DNA Methylation Kit (Zymo Research, Irvine, CA, USA) following the manufacturer’s protocol with the specified modifications for the Illumina Infinium Methylation Assay. After conversion, all bisulfite reactions were cleaned using the Zymo-Spin columns and eluted in 12 μL of Tris buffer. Following elution, BS converted DNA was processed through the Infinium MethylationEPIC array v2.0 protocol (Illumina Inc., San Diego CA, USA). The EPIC array v2.0 contains >930K probes querying methylation sites including CpG islands and non-island regions, RefSeq genes, ENCODE open chromatin, ENCODE transcription factor binding sites, and FANTOM5 enhancers. To perform the assay, 4uL of converted DNA was denatured with 4ul 0.1N sodium hydroxide. DNA was then amplified, hybridized to the EPIC bead chip, and an extension reaction was performed using fluorophore-labeled nucleotides per the manufacturer’s protocol. Array beadchips were scanned on the Illumina iScan platform and probe-specific calls were made using Illumina Genome Studio software. The R package SeSAMe (*70*) was used to process IDAT files generated from the Infinium MethylationEPIC v2.0 array and for downstream differential methylation locus (DML) and differential methylation region (DMR) analysis. Further details on data processing have been provided in supplementary methods.

### Network analysis

Significant genes (p-adj. 0.05) identified from RNA-seq and methylation analysis were analyzed using the STRING database. MCL clustering was used to organize nodes, and the top statistically significant functional enrichment (i.e., lowest FDR value) was displayed above each cluster. TP53 was removed from the gene list because it was identified as a disease-associated variant, rather than a methylation site.

### Statistical Analysis

The alpha-synuclein-SAA data (ThT fluorescence and Fmax) collected from appendices were analyzed using the Mann-Whitney U test. For common proteoforms identified via LC-MS, proteoforms with more than 50% missing values were removed. Raw proteoform counts were log10 transformed, and samples were scaled to 0 mean and a unit standard deviation. Values were then averaged across the replicates. For rare proteoforms, proteoforms with more than 50% missing values were retained. Rare proteoforms were detected if raw counts existed in any of the replicates. For aggregation propensity, proteoform raw counts for each sample were first divided by its total, then multiplied by the aggregation propensity score calculated by PASTA 2.0 (*31*). Values were then averaged across the replicates. We used Limma’s robust linear regression to identify tissue differences, adjusting for disease group and age, and empirical Bayes was used to adjust coefficient estimates. P-values were corrected for multiple testing using FDR. Group differences were tested separately for the appendix and SN. For proteoform abundance, aging difference was tested separately for appendix and SN. Limma’s robust linear regression was fitted on a model that includes group and age interaction. Then contrast.fit with contrast.matrix was used to perform aging difference on control and synucleinopathy. The p-values were corrected for multiple testing using FDR. For aggregation propensity, robust linear regression from Limma was first fitted on a model that includes tissue and age interaction, adjusting for group. Then contrast.fit with contrast.matrix was used to perform aging difference on the appendix and SN. p-values were corrected for multiple testing using FDR.

Common proteoforms were used for clustering analysis. For the remaining proteoforms the missing values were imputed with the MICE package (*71*). Then enhanced hierarchical clustering was performed by R. The clustering result was plotted by factoextra package (*72*). The following tissue and group difference analysis for each cluster was based on the average values of proteoforms inside the cluster. Common proteoforms were used for diagnosis prediction using the R package caret (*73*). Proteoforms that have missing values were imputed using k-nearest neighbor imputation (k=5). The resulting data matrix was used to train a classifier using 10-fold repeated cross-validation (3 repeat iterations). Within each cross-validation step, the data were further preprocessed by removing near-zero variable proteoforms and then scaling and centering each proteoform to a 0 mean and unit standard deviation. In order to assess variable importance, a classifier was trained on all the data using the parameters learned in the cross-validation step.

## List of Supplementary Materials

Supplementary Methods Fig S1 to S4

Data files S1 to S10

## Acknowledgements

We thank the Van Andel Institute’s Pathology and Biorepository, Genomics, Bioinformatics and Biostatistics Cores, and the Michigan State University Genomics Core for assistance with experimentation. We are grateful to the Oregon Brain Bank, Parkinson’s UK Brain Bank, the NIH NeuroBioBank, LifeNet Health and Corewell Health for providing the tissues used in this study. We would like to acknowledge Paul M. Thomas and Eva Boonen for their contributions and support. Finally, we gratefully acknowledge the contributions of the late Dr. Viviane Labrie, whose scientific vision and passion for advancing neurodegenerative disease research significantly shaped this work.

## Funding

This work was supported by the NIH-National Institute of Neurological Disorders and Stroke grant R01NS114409 awarded to LB. The SAA studies were partially funded by U24AG079685 grant to CS and a grant from Michael J. Fox Foundation to SP.

## Author contributions

Conceptualization: PB, LB

Methodology: CS, PB, LB, JHK, RDL

Investigation: EA, SZ, JWS, MS, PL, MP, SP, BAK

Visualization: PL, JWS, JG, MP, SP, BAK

Funding acquisition: PB, LB

Project administration: CS, PB, LB

Supervision: RDL, JHK, SP, CS, PB, LB

Writing – original draft: EA, SZ, JW, BAK

Writing – review & editing: EA, SZ, JW, PB, LB, BAK

## Competing interests

CS is a Founder, Chief Scientific Officer, consultant and shareholder of Amprion Inc., a biotechnology company that focuses on the commercial use of seed amplification assays for high-sensitivity detection of misfolded protein aggregates involved in various neurodegenerative diseases. Sandra Pritzkow also has a conflict in relation to Amprion. The University of Texas Health Science Center at Houston has licensed patents and patent applications to Amprion. During the time that this study was being conducted, PB became an employee of F. Hoffmann-La Roche and obtained stock in the company (one of the data were generated by F. Hoffmann-La Roche). He also has ownership interests in Acousort AB, Axial Therapeutics, Enterin Inc and Kenai Therapeutics.

### Data and materials availability

Custom code for statistical analysis is available at https://github.com/lipeipei0611/SynProteoforms. Due to the current U.S. government shutdown, data deposition in publicly accessible repositories has been temporarily delayed. All datasets supporting the findings of this study will be made publicly available in the designated repositories as soon as repository services are restored. Data are available from the corresponding author upon reasonable request until the repositories reopen.

