## Supplementary Methods for "Aggregation-prone alpha-synuclein proteoforms and dysregulated molecular signatures in the vermiform appendix of synucleinopathy patients"

**Preparation of the tosylactivated magnetic beads**

Preparation of the tosylactivated magnetic beads (Dynabeads MyOne Tosylactivated, Thermo Fisher Scientific) for alpha-synuclein immunoprecipitation was performed according to the manufacturer’s protocol, with minor modifications. Briefly, 500 µL beads were washed with 1 mL coating buffer (0.1 M sodium borate buffer pH 9.5). The beads were resuspended in 715 µL coating buffer with 120 µL of clone 42 anti-alpha-synuclein antibody (BD Transduction Laboratories), followed by addition of 415 µL of an ammonium sulfate solution (3M ammonium sulfate dissolved in coating buffer). The bead mixture was incubated with nutation for 24 h at 37°C. The beads were next collected on a magnet and resuspended in 1.25 mL blocking buffer (PBS pH 7.4, 0.5% BSA w/v, and 0.05% Tween-20), incubating overnight at 37°C with nutation. The beads were then washed 3 times with storage buffer (PBS pH 7.4, 0.1% BSA, 0.05% Tween 20, and 0.02% sodium azide). The beads were suspended in 1 mL storage buffer at 4°C until use.

**Data processing for liquid chromatography-mass spectrometry**

Briefly, a list of chemical formulas for the proteoforms was used to calculate expected isotopic patterns. Then, the centroided peaks of the observed raw spectra above a minimum intensity (NL=100) were searched against the expected m/z of the two most abundant isotopomers calculated for the given chemical formulas. A window of 10 ppm was allowed for matches. Then, the full isotopic distribution for matching formulas was adapted to the observed mass error, and each spectrum was searched for the expected isotopomers with a 2 ppm mass error allowance. The relative intensities of the matched isotopic peaks were compared to the expected distribution in order to generate a fitting score. Fitting scores ranged from 0-1 and were calculated by a least squares model weighed positively towards higher intensities. Proteoform isotopomer intensities were quantified for each scan in which the relative chemical formula passed a fitting score threshold of 0.3. The total proteoform intensity per LC-MS run was calculated for each proteoform by the area under the proteoform-specific chromatographic curve (AUC). The AUC of each form was then normalized by the total ion intensity (or total ion current, TIC) of each LC-MS run. The data were then standardized by calculating standard deviation distance of the normalized AUC of each LC-MS run to the mean of the dataset for each proteoform. These standardized intensities were then compared between samples. Additionally, raw files were processed through TDPortal (<http://nrtdp.northwestern.edu/resource-software>) to identify intact proteins and characterize proteoforms, searching against a highly annotated version of the Human UniProt KnowledgeBase.

**Data processing for RNA sequencing**

Adaptor sequences and low-quality reads were trimmed using Trim Galore (*1*). The trimmed reads were aligned to the GRCh38 reference genome using STAR (*2*) with the parameter “--quantMode GeneCounts”. Differential gene expression (DGE) analysis was performed using DESeq2 (*3*) based on gene counts generated from STAR. In DGE, we first applied function “filterByExpr” from edgeR (*4*) to remove lowly expressed genes (min.count was set to 5). Next, the differential expression analysis between PD and control samples was implemented using DESeq function. Shrinkage of effect size (L), which is useful for reporting and downstream enrichment analysis, was applied using function “lfcShrink”. Differentially expressed (DE) genes were identified using an adjusted p-value cutoff of 0.05. Gene set enrichment analysis (GSEA) was performed using ClusterProfiler (*5*) on Kyoto Encyclopedia of Genes and Genomes (KEGG) and Gene Ontology (GO) terms.

**Data processing for DNA methylation analysis**

The “openSesame” function from SeSAMe was applied to convert IDAT files into DNA methylation level (i.e., beta value) matrices. SeSAMe employs linear models to identify DMLs between Parkinson’s disease (PD) and control groups. For DMR analysis, neighboring CpG sites exhibiting consistent methylation patterns were merged into differentially methylated regions (DMRs), with adjusted p-values (0.05) calculated using the Benjamini-Hochberg procedure. CpG site annotations were extracted from sesameData (*6, 7*), and these annotations were merged and utilized for downstream enrichment analysis. DMRs were designated as CpG islands, CpG shores, CpG shelves, open sea, relative to CpG sites, and as TSS1500, TSS200, 5' UTR, 1st Exon, Body, Intergenic, based on their proximity to nearby genes. “methylRRA” from methylGSA (*8*) was used for enrichment analysis for Encyclopedia of Genes and Genomes (KEGG) and gene ontology (GO). For methylation enrichment analysis, raw p-value from DML was used as input, and hypo- and hyper-methylated CpG sites were analyzed independently. As MethylGSA did not natively support EPIC v2.0 natively at the time of the analysis, custom CpG site annotations (see above) were used for the enrichment analysis.
