## Supplementary figures and images for "Aggregation-prone alpha-synuclein proteoforms and dysregulated molecular signatures in the vermiform appendix of synucleinopathy patients"

### Supplementary Figure 1

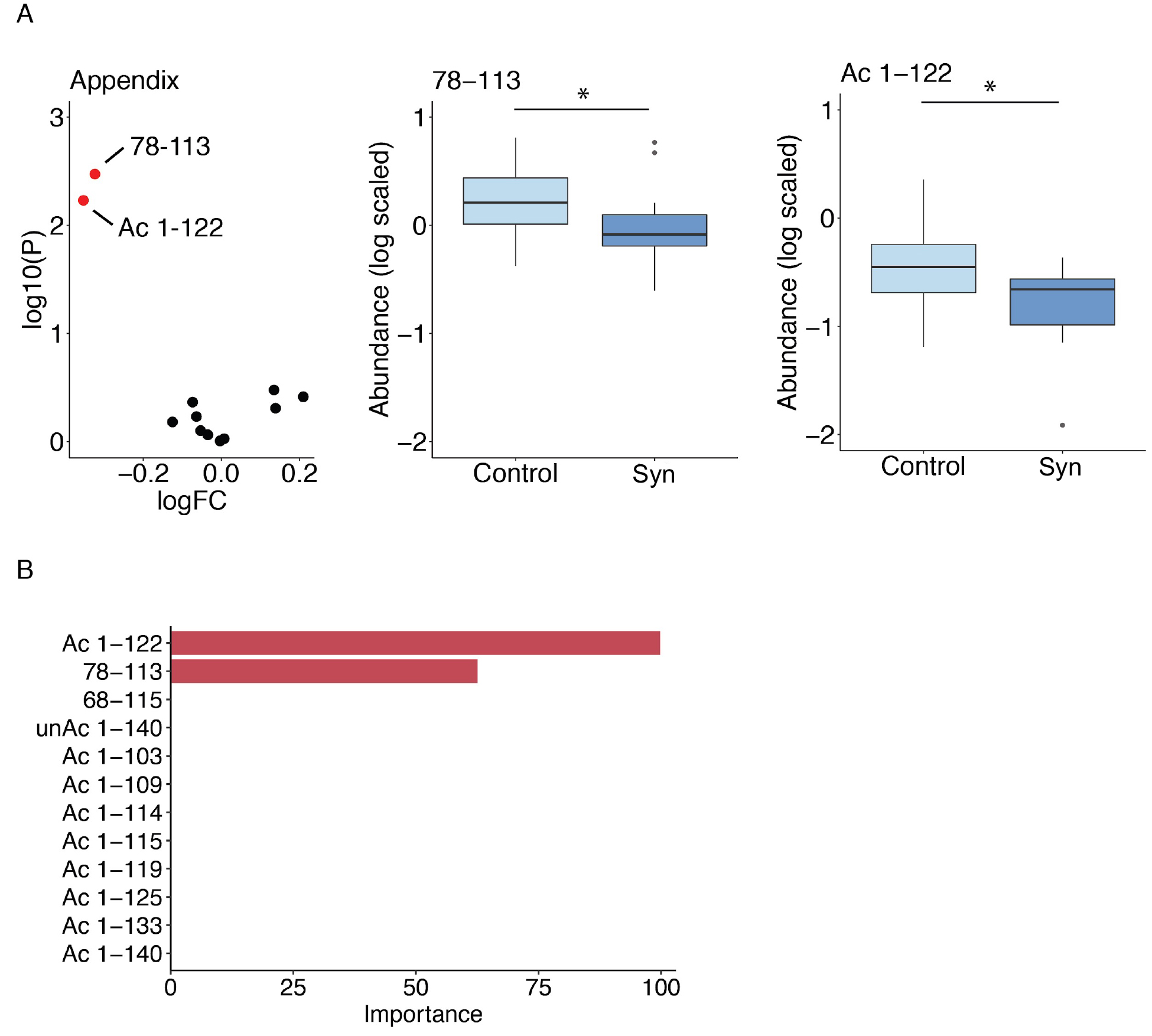

### Supplementary Figure 2

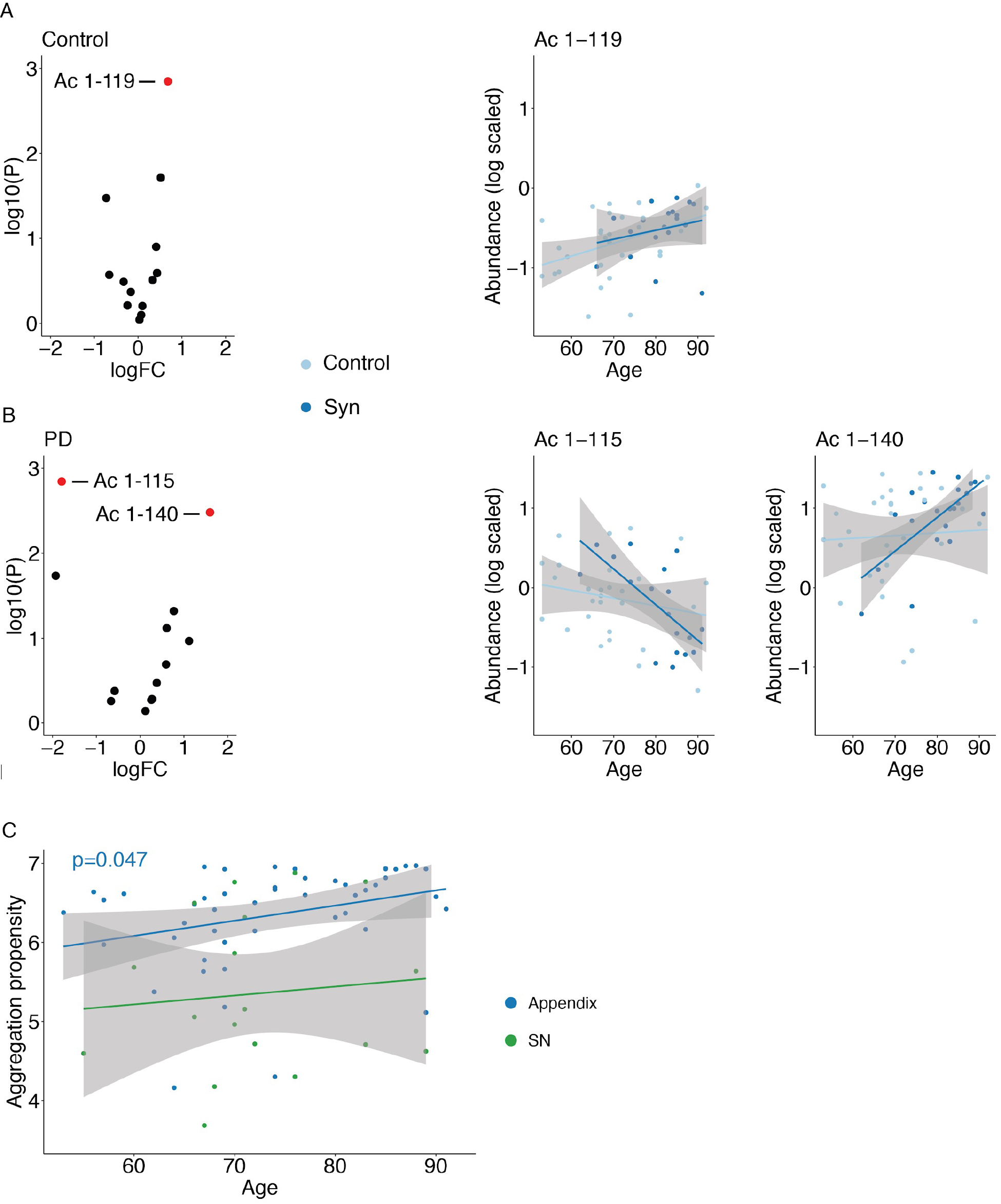

### Supplementary Figure 3

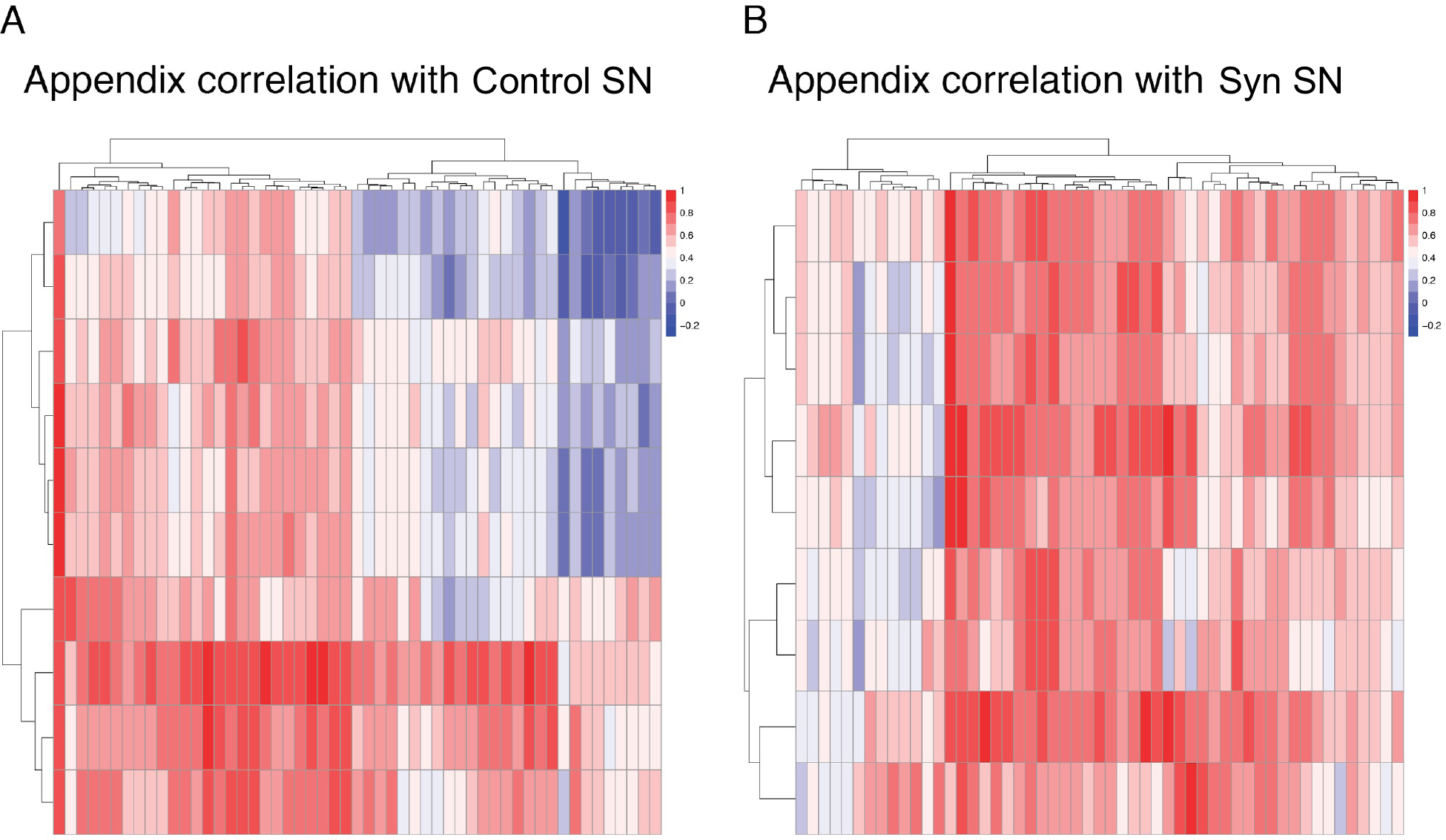

### Supplementary Figure 4

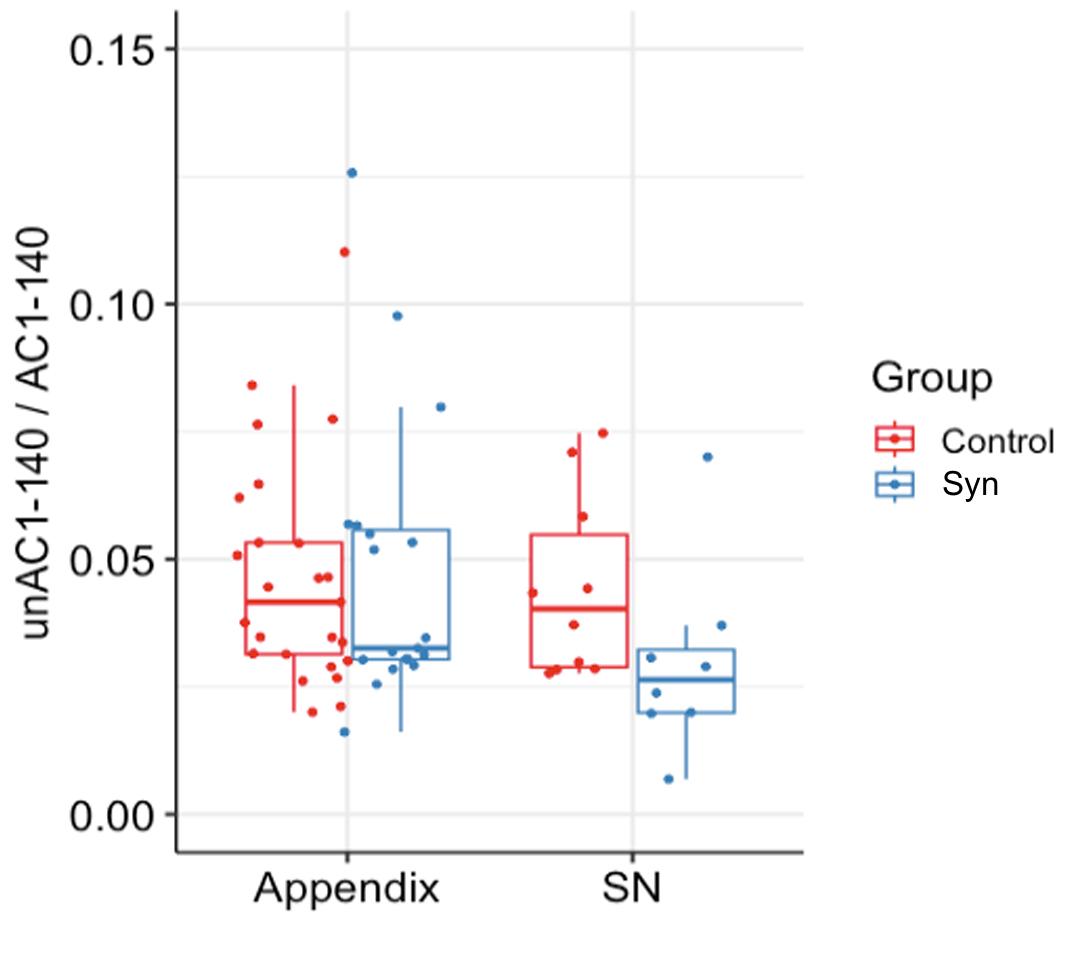
